# An efficient CRISPR-Cas9 enrichment sequencing strategy for characterizing complex and highly duplicated genomic regions. A case study in the *Prunus salicina* LG3-MYB10 genes cluster

**DOI:** 10.1101/2022.01.24.477518

**Authors:** Arnau Fiol, Federico Jurado-Ruiz, Elena López-Girona, Maria José Aranzana

## Abstract

Genome complexity is largely linked to diversification and crop innovation. Examples of regions with duplicated genes with relevant roles in agricultural traits are found in many crops. In both duplicated and non-duplicated genes, much of the variability in agronomic traits is caused by large as well as small and middle scale structural variants (SVs), which highlights the relevance of the identification and characterization of complex variability between genomes for plant breeding. Here we improve and demonstrate the use of CRISPR-Cas9 enrichment combined with long-read sequencing technology to resolve the *MYB10* region in the linkage group 3 (LG3) of Japanese plum (*Prunus salicina*), which has a length from 90 kb to 271 kb according to the *P. salicina* genomes available. We demonstrate the high complexity of this region, with homology levels between Japanese plum varieties comparable to those between *Prunus* species. We cleaved *MYB10* genes in five plum varieties using the Cas9 enzyme guided by a pool of crRNAs. The barcoded fragments were then pooled and sequenced in a single MinION Oxford Nanopore Technologies (ONT) run, yielding 194 Mb of sequence. The enrichment was confirmed by aligning the long reads to the plum reference genomes, with a mean read on-target value of 4.5% and a depth per sample of 11.9x. From the alignment, 3,261 SNPs and 287 SVs were called and phased. A *de novo* assembly was constructed for each variety, which also allowed detection, at the haplotype level, of the variability in this region. CRISPR-Cas9 enrichment is a versatile and powerful tool for long-read targeted sequencing even on highly duplicated and/or polymorphic genomic regions, being especially useful when a reference genome is not available. Potential uses of this methodology as well as its limitations are further discussed.

## BACKGROUND

The origins of agriculture and the development of new crops are tightly linked to whole genome duplications (WGD), and/or to chromosome rearrangements, which are among the major determinants of useful trait diversity (1-3) as well as of genome complexity (4). In addition to WGD, small-scale duplications (SSD), usually produced by segmental duplications and gene expansion, also play an important role in crop innovation (5, 6). Reduced selective pressure acting on duplicated genes may, in some cases, result in them acquiring novel functionalities that contribute to diversification and lead to important traits for agriculture (7-10).

Gene diversification results in high levels of diversity within gene families. For example, a substantial number of copies of celiac disease related α-gliadin genes are estimated in wheat (ranging from 25–35 to 100–150 copies per haploid genome), and are probably generated by duplication, deletion events and retrotransposon insertion (11, 12). These genes show high variability between genomes (13), with at least half of the copies being inactive or pseudogenes (14).

In both duplicated and non-duplicated genes, a large fraction of the variability is caused by large as well as small and middle scale structural variants (SVs). For example, Chawla et al. (15) found that up to 10% of all genes in *Brassica napus* were affected by small- to mid-scale SV events. In tomato, Alonge et al. (16) found multiple SVs, in many cases mediated by transposable elements (TEs); many of the SVs responsible for changes in gene dosage and expression levels modified agronomic traits such as fruit flavor, size, and production. Recent studies of SVs between and within plant species have revealed extensive genome content variation (17), leading to the pan-genome concept, which is centered on the study of the entire gene repertoire of a species by sequencing multiple individuals (18).

The major effect of duplicated genes and SVs, especially in crop agronomic traits, highlights the importance of studying the variability between genomes in these genes for plant breeding.

Whole-genome sequencing (WGS) methods have contributed enormously to broadening our understanding of the genetic basis of useful traits. However, genome complexity still represents a challenge for genome assembly and annotation. In general, large divergent duplications with less than 97% homology can be resolved by WGS assembly, whereas reads of duplications with higher homology are frequently collapsed, producing assembly errors (19).

A large number of genomes have been obtained, and SNPs and small InDels discovered using short-read high throughput sequencing (SR-HTS), but long-read high throughput sequencing (LR-HTS) is required to resolve highly homologous duplicated regions and SVs, including repetitive regions and different forms of copy number variations, such as presence-absence variants (20). Multiple reference genomes for crop species are currently being generated using LR-HTS methods (16, 21, 22); however, although the cost of long-range sequencing methods is decreasing, the availability of large amounts of high quality DNA, whole genome data storage, alignment and computation costs may be drawbacks when the objective is to resolve and explore the variability in a large panel of varieties of a unique complex loci for an interesting agricultural trait.

Over the last few years, several skimming and targeted-enrichment sequencing methods have been released and successfully used to pinpoint small variants in single regions, especially in phylogenomics. However, a cost-effective method to scan the variability (including SV) in a complex or highly diverse region in a panel of genotypes (for example varieties or seedlings) using LR-HTS strategies is still lacking.

Recently, a methodology has been reported for long-read targeted sequencing using CRISPR-Cas9 technology to direct the sequencing adapters to the regions of interest (23, 24). In this procedure, selected regions of high molecular weight dephosphorylated DNA are cut by the Cas9 enzyme directed by specific guide RNAs (gRNAs). The digested fragments are enriched by long-read sequencing thanks to the preferential ligation of sequencing adapters towards the 5’ phosphate groups generated at the cleaved ends. Gilpatrick et al. (23) used a pool of gRNAs to cut and sequence several loci associated with human breast cancer on the same run, resolving SNPs and SVs at the haplotype level. This methodology is simple to perform, as it does not require physical separation of the DNA for sequencing, cloning or amplification steps, thus allowing native DNA strands to be read, and it is not affected by amplification bias and allows visualization of allele-specific methylation patterns (23, 25). CRISPR-Cas9 enrichment has recently been used on a plant genome (as a proof of concept) showing that, at the haplotype level, it can resolve the SV that causes red flesh color in apples (24) Although this methodology was used to sequence a short (7.8 Kb) and low complex region in one apple variety, the successful results augur well for its use for fine-mapping in other situations.

The objective of this study was to evaluate the potential of CRISPR-Cas9 enrichment to identify the variability (in particular SV) in highly complex regions using a pool of DNA of highly diverse varieties. The region selected was the Japanese plum (*Prunus salicina*) *MYB10* region located on linkage group 3 (LG3), found recently to contain at least three *MYB10*.*1* (*MYB10*.*1a, MYB10*.*1b* and *MYB10*.*c*) genes, one *MYB10*.*2* gene and one *MYB10*.*3* gene, while additional copies could not be discarded (26). A marker system designed in this region was able to characterize the LG3-*PsMYB10* allelic organization in Japanese plum varieties and progenies into six haplotypes (H1 to H6), each with a different combination of the *MYB10*.*1–3* alleles (26). For example, H1 and H3 haplotypes contained three *PsMYB10*.*1* alleles, one of them (*MYB10*.*1a*-a356) associated with the anthocyanin skin color; polymorphisms in the promoter and intron of this allele in H1 and H3 were observed, confirming the high variability of this region.

We designed seven gRNAs in conserved sites of the *PsMYB10* genes and pooled them, together with two guides flanking the region, to allow for multiple Cas9-mediated cuts in the DNA of five Japanese plum commercial varieties, all with a distinct and heterozygous genotype for the *MYB10* region, while sharing one haplotype by pairs. The fragments were labeled with a barcode system and pooled and sequenced in a single MinION Oxford Nanopore Technology (ONT) run. Reads alignment with reference genomes or *de novo* alignments successfully identified SNPs and SVs, most of them newly discovered and others observed in our previous work, so validating the methodology for sequencing and variant detection of regions containing complex duplications.

## RESULTS

### The Japanese plum LG3-MYB10 region is highly complex

We used the Japanese plum LG3-MYB10 region as a model of high complexity. Japanese plum is a self-incompatible fruit tree with a complex history of interspecific crosses between diploid plums: the LG3-MYB10 region in *Prunus* genomes, particularly in Japanese plum, is highly diverse and contains a cluster of at least three paralogous *MYB10* gene copies.

To test the complexity of this region and validate its suitability for analysis, we identified and compared the assembly of the LG3-MYB10 region in the two Japanese plum genomes currently available (one for ‘Sanyueli’(27) and one for ‘Zhongli No. 6’ (28) varieties). In the ‘Sanyueli’ genome (v1.0) the region spanned 135 kb in chromosome 4 (Chr4:12192580-12327184). Although the genome annotation contained only one gene, homologous to *PsMYB10*.*2*, BLAST identified an additional *PsMYB10*.*2*, two *PsMYB10*.*1* and one *PsMYB10*.*3* gene copies in the region.

In the ‘Zhongli No. 6’ genome we identified two MYB10 regions 2 Mb apart in chromosome 3: one 271 kb long (from now on, Zhongli-1; LG03:30838572-31109359) and the other 90 kb long (Zhongli-2; LG03:28592935-28682529). The Zhongli-1 region had five *MYB10* genes annotated, corresponding to two copies of *MYB10*.*1*, two copies of *MYB10*.*2* and one copy of a *MYB10*.*3* gene. These genes were mis-annotated: the second exon was omitted in all but one of the *PsMYB10*.*1* copies, while in this copy the STOP codon in the last exon was not detected, causing the spurious automatic identification of five extra exons along the next 6 kb of sequence. By BLAST we identified three additional genes: one was homologous to *MYB10*.*1* and two homologous to *MYB10*.*2*. In the Zhongli-2 region, four *MYB10* genes were annotated: two *MYB10*.*1* copies, one *MYB10*.*2* and one *MYB10*.*3*. The second exon was also mis-annotated in most of the gene copies of this region. BLAST analysis did not identify additional *MYB10* genes.

To identify and visualize the differences in size and gene copy number and organization, we produced dot plots comparing the three LG3-MYB10 region assemblies (**Figure 1**). These reflected the high variability and poor homology along the region, with inverted assemblies, mis-assemblies and missing fragments. The Zhongli-1 MYB10 region was assembled in an inverted orientation, when compared with ‘Sanyueli’ and Zhongli-2. Despite this, the Zhongli-1 (of 271 kb) and the Zhongli-2 (90 kb) alignments were collinear and with high homology (69% of sequence hits), with an evident gap of ∼150 kb due to a missing fragment in Zhongli-2 explaining most of their difference in size (270 kb vs 90 kb). The percentage of hits between ‘Sanyueli’ and Zhongli-1 was lower (51%), with multiple misalignments along the region that explained the smaller size in ‘Sanyueli’. Similarly, a discontinuous alignment and an inversion was also observed between Zhongli-2 and ‘Sanyueli’ (51% of sequence hits).

**Figure 1.**
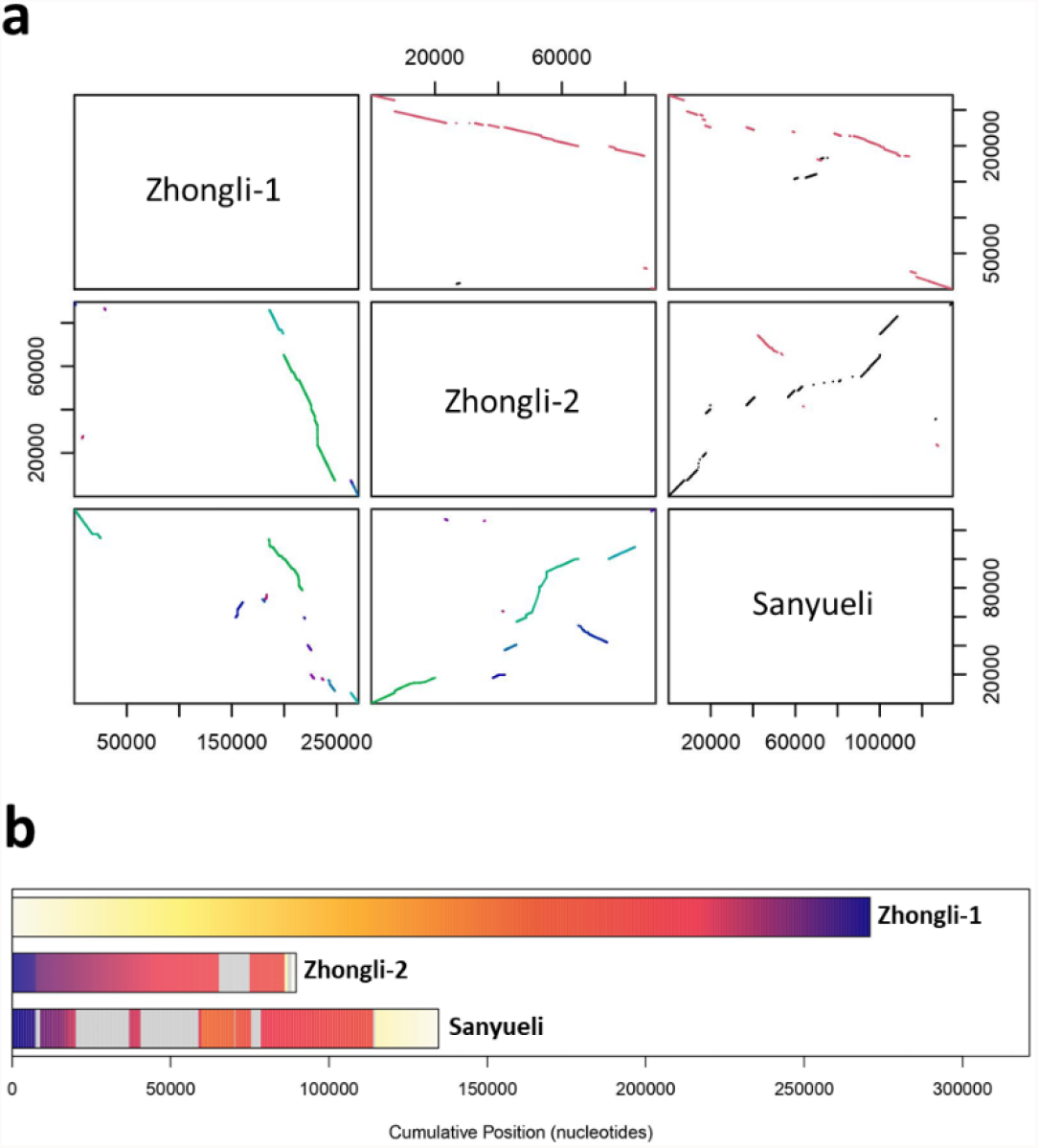
**a** Dot plot comparison of the three LG3-MYB10 assemblies in the two Japanese plum genomes published: ‘Zhongli No. 6’ (with two assemblies: Zhongli-1 and Zhongli-2) and ‘Sanyueli’. Top right of the diagonal: homologous hits; inversions colored in red. Bottom left of the diagonal: homologous blocks colored by their score, from green to colder colors while the homology value decreases. Coordinates of each region are written in base pairs for both axes. **b** Distribution of the homologous sequences in the three compared genome regions. Grey areas correspond to sequences not found in the Zhongli-1 MYB10 region. Homologous regions are represented in the same yellow to purple colors.

Similar comparisons were constructed for other *Prunus* species with more than one available reference genome (**Additional File 1, Additional File 2**). The MYB10 regions of the two sweet cherry (*P. avium*) genomes were highly collinear with 75% of sequence hits. In peach (*P. persica*), the region was collinear and highly similar in the ‘Lovell’ and ‘Chinese Cling’ genomes (85% of sequence hits), and three times larger than in the wild relative *P. mira*, with a mean of 55.5% of sequence hits. In apricot, we compared five *P. armeniaca* accessions and three wild apricot species (two *P. sibirica* and one *P. mandshurica*). Homology between each *P. armeniaca* pairwise comparison ranged from 44% to 91% (mean of 67.7% of hits). When these were compared to the wild apricot genomes, the homology ranged from 58% to 93% (mean of 74.8% of hits) with *P. sibirica*, and from 47% to 59% (mean of 53.2% of hits) with the *P. mandshurica* genome. The mean percentage of sequence hits between the *Prunus* sections was 19.4% with values ranging from 8.6% to 37%.

### The designed gRNAs successfully cut LG3-MYB10 genes

For selective high-throughput sequencing of the desired LG3-MYB10 region in Japanese plum varieties, we designed seven CRISPR RNAs (crRNAs) in genic regions (exons and introns) of the *PsMYB10*.*1, PsMYB10*.*2* and *PsMYB10*.*3* genes, including regions conserved between the genes to minimize the number of required crRNAs. The crRNAs were designed at the two DNA strands to ensure forward and reverse sequences and to direct the sequencing from the inside to the outside of the gene, covering the whole region with concatenated sequences (**Figure 2**). The *PsMYB10*.*1, PsMYB10*.*2* and *PsMYB10*.*3* sequences obtained in Fiol et al. (2021), the peach ‘Lovell’ reference genome, and other *Prunus* genomes were used to design the crRNAs with higher on-target capacity and lower off-target risk (see Methods). Therefore, crRNA-s1 was designed to cut within the exon 2 of *MYB10*.*1* and *MYB10*.*2* genes in the plus strand, while crRNA-s3 was directed to the same exon and strand in *MYB10*.*3* (**Additional File 3**). To avoid a putative mis-cut due to a possible SNP in the second exon of the *MYB10*.*2* cleavage site of crRNA-s1 (although only identified in *P. cerasifera, P. domestica* and *P. avium*) we designed crRNA-s2 for the alternative nucleotide. As reported by Fiol et al. (2021), variability in the introns was higher than in exons, so the crRNAs were designed taking into consideration the variants known. Introns of *PsMYB10*.*1, PsMYB10*.*2* and *PsMYB10*.*3* were targeted with crRNA-a1, crRNA-a3 and crRNA-a4, respectively, while crRNA-a2 was designed to include an SNP variant in one of the *PsMYB10*.*1* alleles present in the sample (*PsMYB10*.*1*-H1 in (26)). In addition, to cut the MYB10 genomic region at each flank, two crRNAs (crRNA-f1 and crRNA-f2) were designed in conserved exon sequences of the nearest flanking genes upstream (Prupe.3G162900 in peach, Pav_sc0000464.1_g320.1.mk in sweet cherry genomes) and downstream (Prupe.3G163400 in peach, Pav_sc0000464.1_g090.1.mk in sweet cherry genomes). (**Figure 2, Additional File 3**).

**Figure 2.**
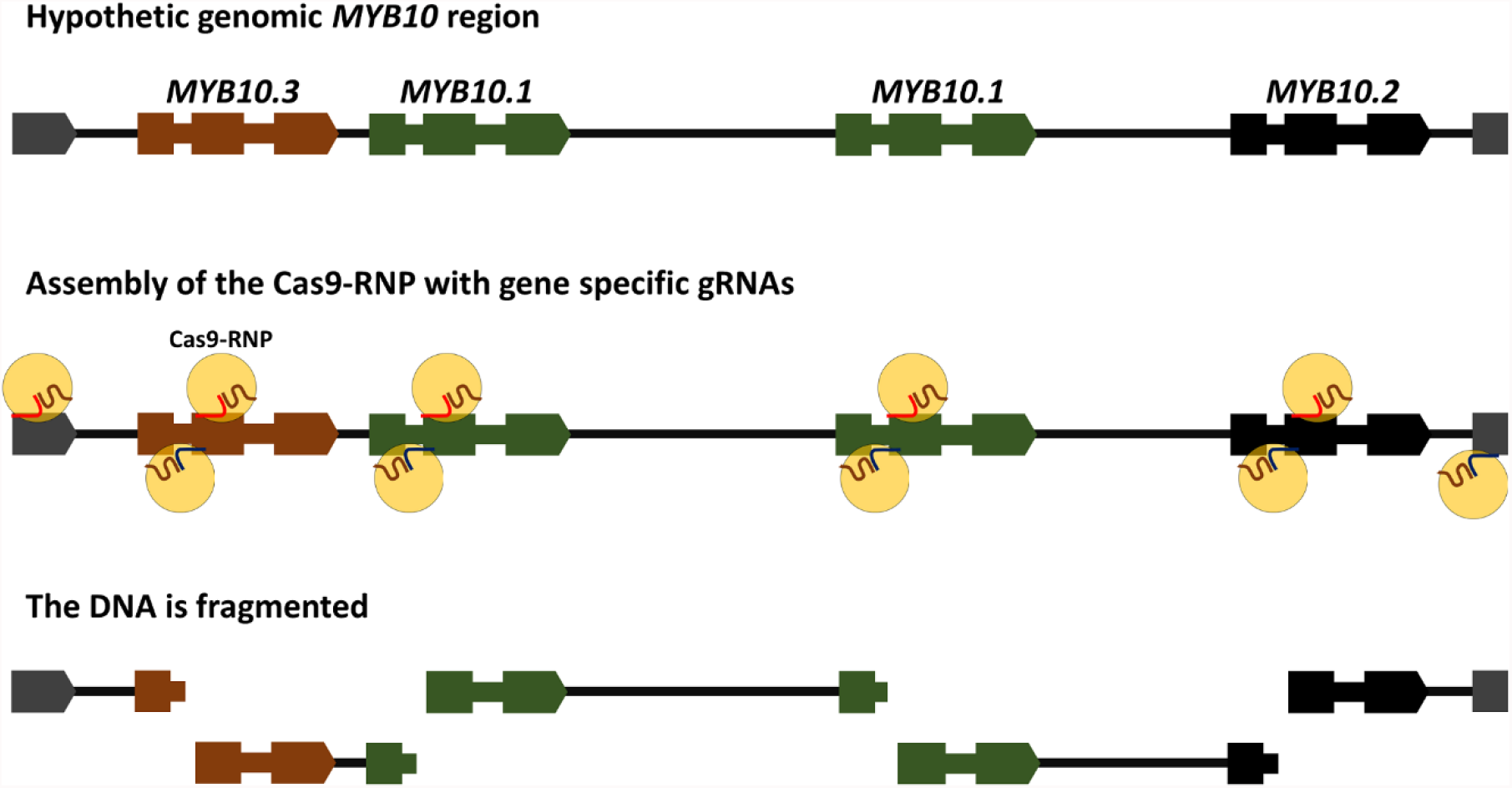
Schematic representation of the CRISPR RNA (crRNA) design to specifically cleave and sequence a hypothetic LG3-MYB10 genomic region in Japanese plum. The guide RNAs (gRNAs) are formed by the interaction of trans-activating RNAs with crRNAs, then with the Cas9 enzyme are assembled into the Cas9-Ribonucleoprotein (Cas9-RNP) complex. The Cas9-RNPs allows the specific cleavage of the LG3-MYB10 region, generating cuts at the two DNA strands for further sequencing in both directions from the inside to the outside of the genes. Gene number, order and size of the region, as well as the expected number of fragments were unknown at the time of the crRNA design.

We first validated the cleavage ability of the crRNAs in PCR amplified DNA. For that we amplified the *MYB10*.*1, MYB10*.*2* and *MYB10*.*3* complete genes in the plum variety ‘Angeleno’ and cleaved the products with a pool of the seven gRNAs targeting the *MYB10* genes only. As intron 2 in one of the *PsMYB10*.*1* alleles in ‘Angeleno’ is about 1.5 kb larger (26), the *MYB10*.*1* primers yielded fragments visualized in two bands in **Figure 3a** (the one containing the large intron is very faint in the gel, due to the preferential amplification of the shorter one). The fragments were cut and visualized in five bands. Three of them corresponded to the fragments 2 (526 bp), 3 (1078 bp), 4 (613 bp) and 5 (991 bp) in the *in-silico* design shown in **Figure 3b**. The two faint additional bands corresponded to fragments 3 and 5 (of about 2.5 kb) of the allele with the large intron 2, and to an undigested fraction of the product of the preferentially amplified copy. Similar results were observed for the *MYB10*.*2* amplicon, which was cut into four fragments. The band corresponding to the small gene fragment (fragment 6 in **Figure 3b**) spanning the exon and intron crRNA sites was not produced in either the *MYB10*.*1* or the *MYB10*.*2* cleavages. The pool was not able to cut the *MYB10*.*3* amplicon. A posterior sequencing of that amplicon revealed an unexpected 210-bp deletion affecting part of intron 1 and all of exon 2, and so lacking the two scission target sites.

**Figure 3.**
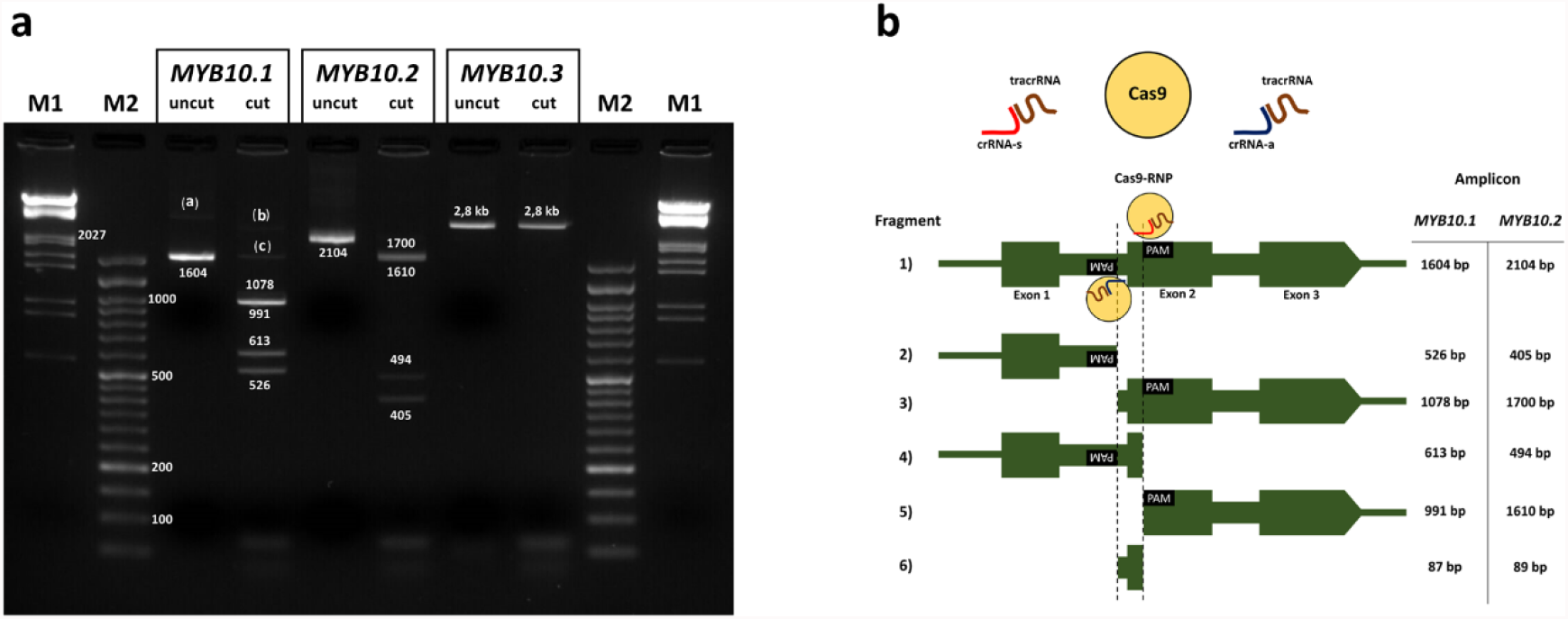
**a** MYB10 amplicon bands before and after cleavage with a pool of the guide RNAs (gRNAs) targeting the genes, with their size annotated in base pairs (bp). Bands (a, b) correspond to the faint bands from the allele with a large intron 1, (c) to the remaining undigested product. Wells M1: Lambda DNA HindIII-EcoRI digested molecular weight marker; M2: DNA Ladder 50 bp ready-to-use (GeneON). **b** Expected fragment sizes after *MYB10*.*1* and *MYB10*.*2* digestion with the Cas9-Ribonucleoprotein (RNP) complex. Guide-RNAs assembled with the trans-activating CRISPR RNA (tracrRNA) and crRNA-s (s1, s2 or s3) cut the exon 2 of the gene, gRNAs with crRNA-a (a1, a2, a3 or a4) cut by intron 1. Fragment 6 was not obtained after cleavage, fragment 3 and 5 bands overlapped in the gel.

### Sequencing, alignment to the reference genomes and variant calling

The genomic DNA of five plum varieties (**Table 1**) was cleaved with the pool of gRNAs and barcoded. A pool of the output reactions was sequenced with the ONT MinION device, which produced 194 Mb of sequences in 33,794 reads of average mean length of 6,097 bp (**Table 1**). After demultiplexing the reads into varieties through their barcode, we obtained sequence yields ranging from 29.32 Mb to 45.64 Mb in reads of mean length from 4,110 to 7,778 bp; the average N50 was 14,510 bp.

**Table 1.** Sequencing statistics for each plum variety after read demultiplexing.

| Variety | Barcode | Total reads | Yield (Mb) | Mean length (bp) | N50 | Mean Q |
| --- | --- | --- | --- | --- | --- | --- |
| ‘Angeleno’ | NB01 | 4,995 | 38.85 | 7,777.9 | 19,049 | 11.0 |
| ‘Black Gold’ | NB02 | 11,106 | 45.64 | 4,109.7 | 10,288 | 10.9 |
| ‘Fortune’ | NB03 | 5,417 | 29.32 | 5,412.2 | 11,786 | 10.8 |
| ‘Golden Japan’ | NB04 | 5,063 | 34.98 | 6,908.0 | 16,740 | 11.0 |
| ‘TC Sun’ | NB05 | 7,213 | 45.27 | 6,275.8 | 14,686 | 10.9 |

The reads of each of the five varieties were aligned against the ‘Sanyueli’ and ‘Zhongli No. 6’ genomes. The coverage and depth of the alignments within the MYB10 region varied for each sample and reference genome used **(Table 2, Additional File 4**). Higher coverage was observed for the alignments against the shorter region of Zhongli-2 (90 kb long, 63.4% of coverage in average) while the coverage of the alignments against Zhongli-1 (54.2% of coverage, 271 kb long) was only slightly lower than that for ‘Sanyueli’ (57% of coverage, 135 kb long). The on-target alignment did not correlate with the number of reads yielded, which was uneven for each variety. In particular, ‘Golden Japan’ was the variety with the lowest number of on-target aligned reads (168, 67 and 45 in ‘Sanyueli’, Zhongli-1 and Zhongli-2, respectively). The mean sequence depth was also unequal for each variety-reference genome combination. In general, for all varieties, a higher depth and higher number of on-target aligned reads was observed with ‘Sanyueli’ **(Table 2)**. The variety with the lowest depth was ‘Golden Japan’ due to the poor alignment, while ‘TC Sun’ performed better.

**Table 2.**
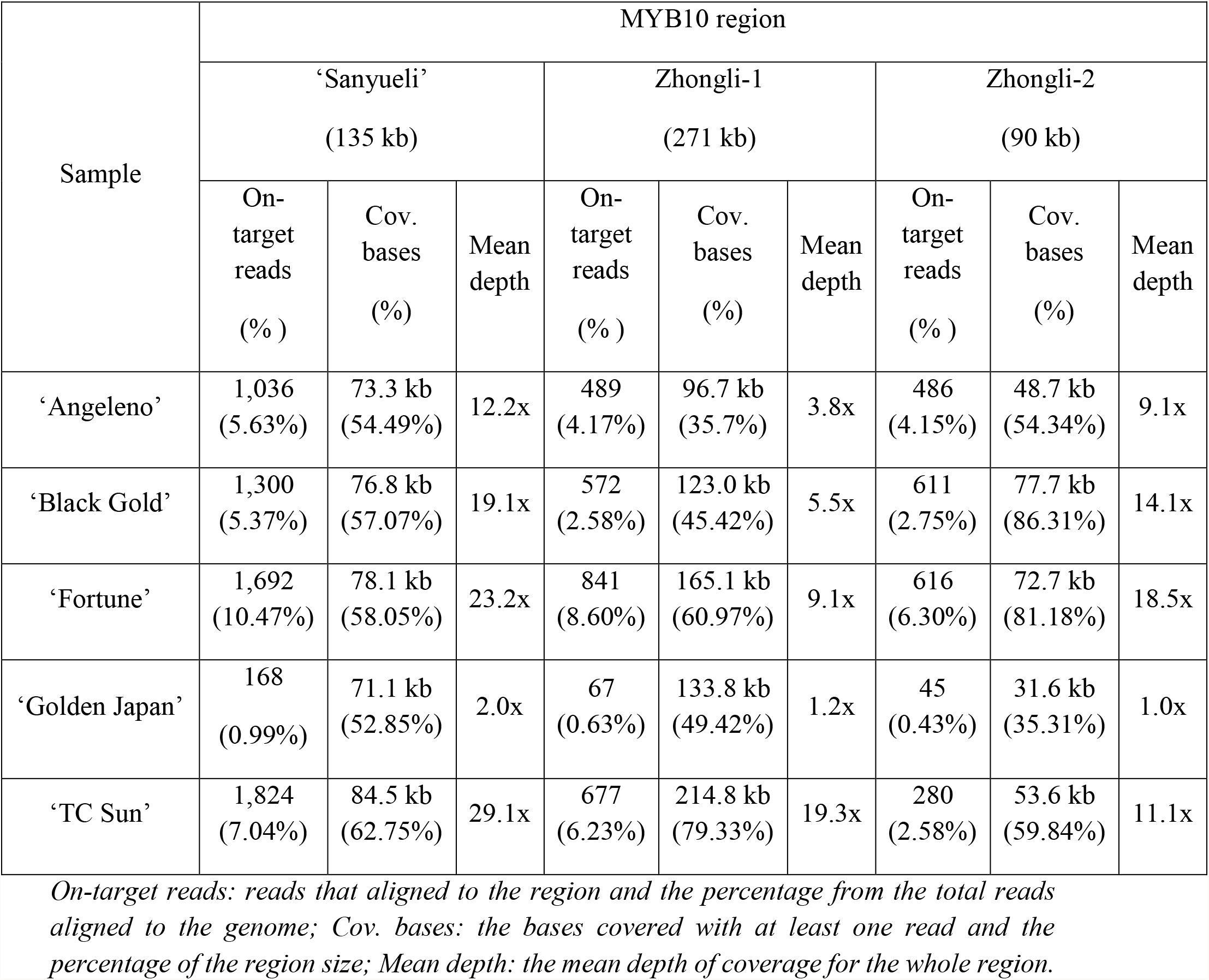
Alignment of each cultivar to the three Japanese plum MYB10 reference regions.

| Sample | MYB10 region |  |  |  |  |  |  |  |  |
| --- | --- | --- | --- | --- | --- | --- | --- | --- | --- |
|  | ‘Sanyueli’<br>(135 kb) |  |  | Zhongli-1<br>(271 kb) |  |  | Zhongli-2<br>(90 kb) |  |  |
|  | On-target<br>reads<br>(%) | Cov.<br>bases<br>(%) | Mean<br>depth | On-target<br>reads<br>(%) | Cov.<br>bases<br>(%) | Mean<br>depth | On-target<br>reads<br>(%) | Cov.<br>bases<br>(%) | Mean<br>depth |
| ‘Angeleno’ | 1,036<br>(5.63%) | 73.3 kb<br>(54.49%) | 12.2x | 489<br>(4.17%) | 96.7 kb<br>(35.7%) | 3.8x | 486<br>(4.15%) | 48.7 kb<br>(54.34%) | 9.1x |
| ‘Black Gold’ | 1,300<br>(5.37%) | 76.8 kb<br>(57.07%) | 19.1x | 572<br>(2.58%) | 123.0 kb<br>(45.42%) | 5.5x | 611<br>(2.75%) | 77.7 kb<br>(86.31%) | 14.1x |
| ‘Fortune’ | 1,692<br>(10.47%) | 78.1 kb<br>(58.05%) | 23.2x | 841<br>(8.60%) | 165.1 kb<br>(60.97%) | 9.1x | 616<br>(6.30%) | 72.7 kb<br>(81.18%) | 18.5x |
| ‘Golden Japan’ | 168<br>(0.99%) | 71.1 kb<br>(52.85%) | 2.0x | 67<br>(0.63%) | 133.8 kb<br>(49.42%) | 1.2x | 45<br>(0.43%) | 31.6 kb<br>(35.31%) | 1.0x |
| ‘TC Sun’ | 1,824<br>(7.04%) | 84.5 kb<br>(62.75%) | 29.1x | 677<br>(6.23%) | 214.8 kb<br>(79.33%) | 19.3x | 280<br>(2.58%) | 53.6 kb<br>(59.84%) | 11.1x |

*On-target reads: reads that aligned to the region and the percentage from the total reads aligned to the genome; Cov. bases: the bases covered with at least one read and the percentage of the region size; Mean depth: the mean depth of coverage for the whole region.*

In general, the depth along the region was unevenly distributed, with scattered regions of high depth (**Additional File 4)**. This was caused by sequences that aligned partially and were clipped from one or both sides, indicating low homology with the genomes used as reference. Sequence clipping was not observed in ‘TC Sun’ aligned to Zhongli-1.

Variants were called for the alignment with ‘Sanyueli’, the one with the highest depth. A total of 3,261 SNPs in the MYB10 region were found, 62.34% of them (2,033) being present in at least two varieties (**Figure 4a**), 98.5% with the same alternative allele (**Additional File 5**). All six observed MYB10 haplotypes (H1–H6) defined by Fiol et al. (2021) plus one inferred (H9) were present in the five varieties, three of them shared by pairs (H1 by ‘Angeleno’ and ‘Black Gold’; H3 by ‘Angeleno’ and ‘Fortune’; and H4 by ‘TC Sun’ and ‘Golden Japan’). This was used to assign the phase of some of the polymorphisms: ‘Angeleno’ and ‘Black Gold’ had 259 private SNPs which may correspond to the shared H1; ‘Angeleno’ and ‘Fortune’ had 62 that could correspond to the shared H3. ‘Fortune’ and ‘Black Gold’ had 87 private SNPs which should be in H2 and H6. Unfortunately, due to the poor quality of the ‘Golden Japan’ sequencing the phase of the SNPs in H4, H5 and H9 could not be inferred.

**Figure 4.**
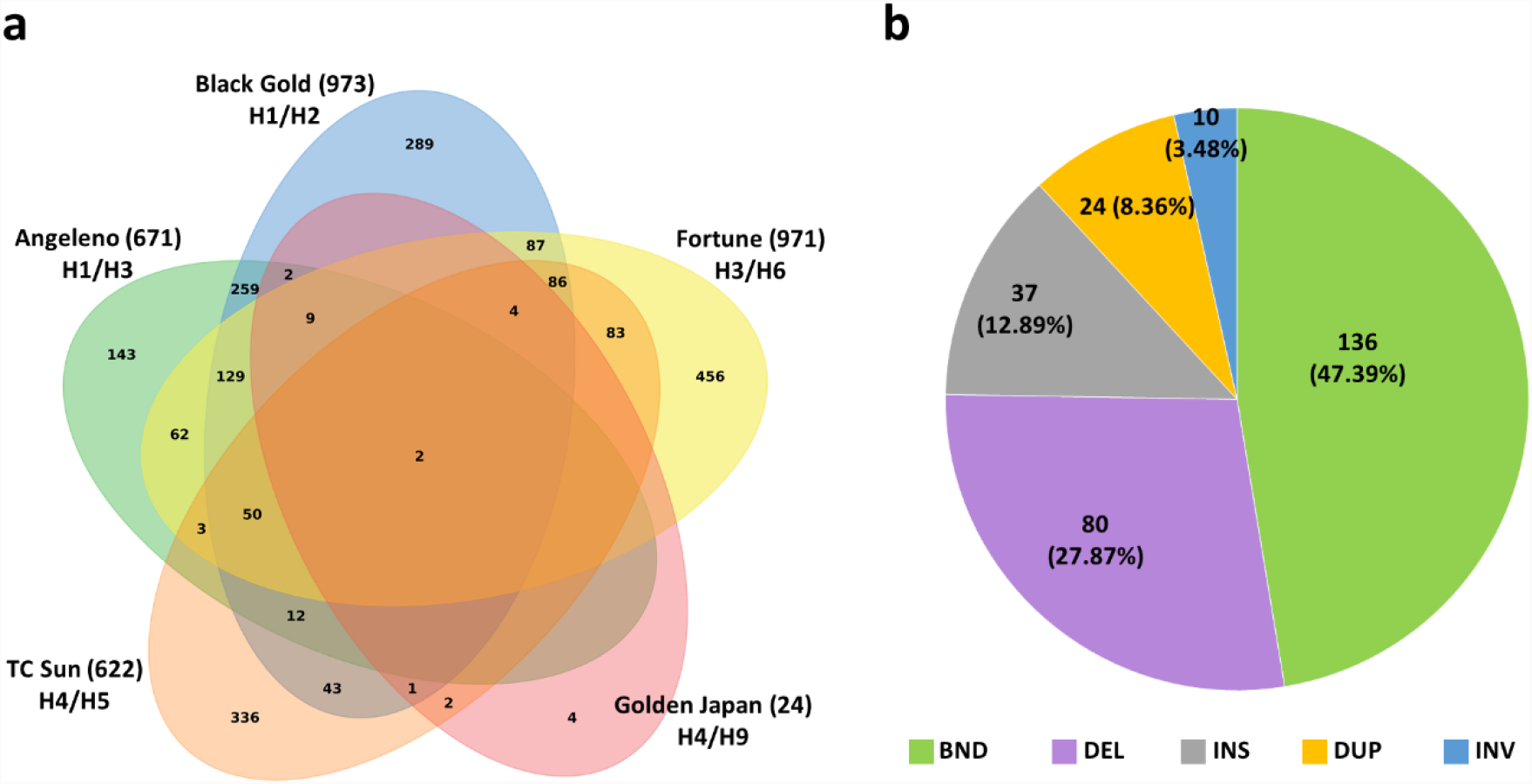
**a** Venn diagram showing the number of SNPs identified in the MYB10 region from each of the five Japanese plum samples sequenced and the overlap between them. The number next to the cultivar stands for their total SNP count. **b** Classification and percentage of all the structural variants identified: breakends (BND), deletions (DEL), duplications (DUP) and inversions (INV).

Two different software tools were used to call SVs and polymorphisms affecting more than 30 bps, identifying 287 variants: 72 in ‘Angeleno’, 63 in ‘Black Gold’, 68 in ‘Fortune’, 30 in ‘Golden Japan’ and 54 in ‘TC Sun’. Most corresponded to sequence breakends (47.39%), followed by deletions (27.87%) and insertions (12.89%) (**Figure 4b**).

To check the accuracy of the SVs call we searched for the 44-bp insertion containing a G-box motif in the *PsMYB10*.*1a* promoter in the H1, H3 and H9 haplotypes in Fiol et al. (2021). The insertion was successfully annotated in ‘Angeleno’ (H1/H3), ‘Black Gold’ (H1/H2) and ‘Golden Japan ‘(H4/H9), but not in ‘Fortune’ (H3/H6).

### *de novo* assembly and homology visualization

Despite being able to identify new polymorphisms with this method, we carried out a *de novo* assembly to overcome the low coverage caused by sequence clipping in the regions with low homology. First we constructed an assembly for each variety, then we used the ‘Zhongli No. 6’ and ‘Sanyueli’ genomes to scaffold the contigs along the region. As expected, the degree of homology and collinearity varied for each assembly (**Additional File 6)**. In general, higher homology was observed in comparisons of the assemblies of the five varieties sequenced with the CRISPR-Cas9 selective cleavage strategy, particularly in those sharing a haplotype. In particular, the assemblies of ‘Angeleno’ and ‘Black Gold’ were always the most similar, with hits ranging from 41% to 53%, depending on the reference region used for assembly.

### Identification of polymorphisms in the *PsMYB10*.*1a* promoter

Given the low homology between the samples and the ‘Zhongli No. 6’ and ‘Sanyueli’ genomes, we searched for polymorphisms among the *de novo* assemblies only. Using as an example a 250 bp contig upstream *PsMYB10*.*1*, previously sequenced with Sanger by Fiol et al. (26), we successfully identified the SNPs and InDels in heterozygosis, among them the 41 bp deletion in H3 and the 44 bp insertion in the H1, H3 and H9 haplotypes. The former (position 66 to 76 in the contig, **Figure 5**) was shared by ‘Angeleno’ (H1/H3) and ‘Fortune’ (H3/H6). The latter (position 200 to 243 in the alignment, **Figure 5**) was present in the varieties with H1, H3 or H9, in agreement with the results of Fiol et al. (26). Most of the SNPs detected were consistent with those previously found, however we identified a few sequencing errors. This was the case of the SNPs in positions 52, 137, 175, and 244–245 in the alignment (**Figure 5**).

**Figure 5.**
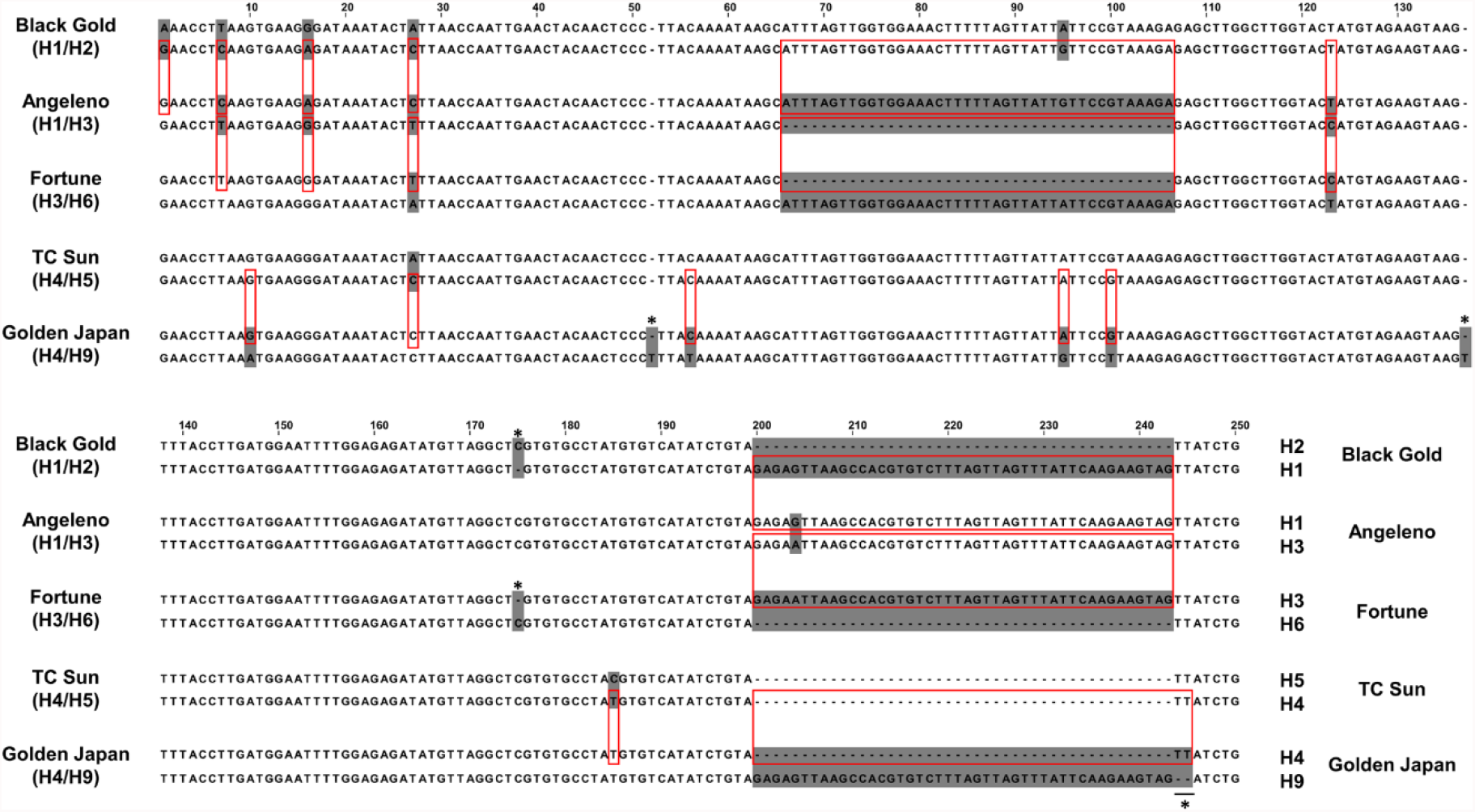
Sequence comparison of the five Japanese plum varieties sequenced. Heterozygous positions (shadowed bases) matched the position on the same haplotype in a different variety (red rectangles). Sequencing mistakes are marked with asterisks.

Despite these few sequencing errors, intrinsic to the ONT sequencing technology, the results validate this methodology for the resolution and identification of variability in highly complex regions.

## DISCUSSION

In this work we confirm sequence enrichment CRISPR-Cas9 selective cleavage as a cheap and efficient high throughput strategy to identify variants and validate its use in highly complex regions in heterozygous genomes. Complex genomic regions in plant genomes are the basis of agriculture and crop diversity. They frequently include small and large variants, transposable elements (TE) and clusters of tandem or segmental duplicated multi-copy genes. Such genes are more likely to carry higher levels of variation (29) and be affected by presence/absence variation (PAV) than singleton genes (30), and often intervene in processes related to the evolution of agronomic traits such as secondary metabolisms, response to environmental stimuli, plant defense response and stress tolerance, flowering time, grain size and fruit traits (29, 31-34). This makes characterizing such regions highly relevant for evolutionary studies as well as for plant breeding (35).

It is difficult, or even impossible, to completely resolve plant genome complexity by short read sequencing. Long-read approaches have improved the contiguity of genome assemblies, while special effort is currently being made to resolve particularly complicated regions of relatively small sizes in selected genotypes. In humans, several authors have used a strategy based on enrichment and sequencing using targeted cleavage of chromosomal DNA with Cas9 combined with Nanopore sequencing to resolve genomic SVs as well as mobile elements (23, 36-38). In plants, this strategy has been validated to fine map short regions (24) and to characterize the transposon insertion landscape (39). Here we extend this strategy for the characterization of much longer regions in a pool of genotypes in species without a reference genome or in regions with large variability in gene content.

As the region of interest (ROI), we selected the *Prunus* chromosome 3 *MYB10* region, which consists of a cluster of *MYB10* gene copies with a gene content variable among *Prunus* species. The genotypes studied were five commercial varieties of Japanese plum. Commercial varieties of this fruit crop derive from multiples crosses of *P. salicina* with other diploid plum species, which enhanced thefruit quality (40).

To reflect the complexity within the ROI we analyzed the synteny within and between *Prunus* sections, including two recent *P. salicina* genomes (one for the variety ‘Sanyueli’ and one for ‘Zhongli No. 6’) that were released when our study was already at a very advanced stage. Despite belonging to the same species, the level of homology in this region of these two Japanese plum varieties was low, comparable to that observed between cultivated and wild members from the same section of apricot and peach. By contrast, the homology within each of the *Prunus* sections studied (apricot, sweet cherries and peach) was higher or moderately higher than those obtained for Japanese plums. Such low level of homology in the ROI within *P. salicina* could be attributable to a possible interspecific origin of the varieties together with a putative mis-assembly of the region in the ‘Zhongli No. 6’ genome, in which two LG3-MYB10 regions were assembled 2 Mb apart instead of the single one expected for *Prunus*. The low identity between the *Prunus* ROI contrasts with the high synteny described at the whole genome level within the *Prunus* genus (41) and confirms the high complexity of this region in *Prunus* in general and in Japanese plums in particular, which validates its selection as a ROI for this study.

Previously in Fiol et al. (26), we observed six haplotypes from the segregation of *PsMYB10* alleles in six F1 Japanese plum progenies and advanced breeding lines, finding high allelic variability and the triplication of the *MYB10*.*1* gene in some haplotypes. Using the limited homology between *Prunus* genomes and by designing numerous PCR primer combinations, we were able to clone some of the alleles and phase the variants. This was a slow and costly strategy, and we were not able to obtain the full sequence of the haplotypes.

In contrast, here we complemented the Cas9 enrichment strategy with a multiplex approach to decrease the cost of sequencing six of the observed haplotypes (named H1 to H6). All the crRNAs were pooled for cleavage of the DNA in a single reaction, and the barcoded fragments of each variety were pooled, in equimolar proportions, for their sequencing on a MinION device. The five varieties selected carried the haplotypes in heterozygosis, three of them (H1, H3 and H4) shared by pairs. While this design was useful for optimizing the sequencing sample set and for phasing the variants, it was not determinant for the success of the methodology.

The crRNAs were designed within conserved regions of intron 1 and exon 2 of the *Prunus MYB10* genes. The *MYB* genes encode a family of transcription factors (TF) present in all eukaryotic organisms. The MYB10 proteins belong to one of the largest TF groups, the R2R3-MYB class, one of the four *MYB* gene classes in seed plants (42). As occurs with the majority of multi-copy gene families, multiple phylogeny and diversity studies in plant lineages have identified conserved motifs within the gene sequence. Such regions can be successfully used to direct crRNAs.

### Design of the crRNAs

Here we targeted some of the conserved regions identified in Fiol et al. (26) in addition to two regions, one in each of the two closest genes up and downstream of the *MYB10* cluster, to border the ROI. This work was started several months before the two Japanese plum genomes were released. Although their availability could have helped in designing the crRNAs, our results indicate that this strategy can be successfully applied to those species lacking a reference genome.

To obtain the sequences of the genes and their upstream and downstream regions, we designed seven non-overlapping forward and reverse crRNAs able to direct the sequencing in both directions. Due to the small length of the *MYB10* sequences available and the high variability in the region, which limited the target region, the on-target activity of the crRNAs was far from optimal (51 to 68, on a scale of 1–100). Their cleavage activity was tested *in vitro* with *MYB10* amplicons of the variety ‘Angeleno’. The results revealed that while the activity values would likely not perform well for *in vivo* gene editing, they were sufficient for *in vitro* cleavage with the system used here. In addition, *in vitro* digestion revealed that fragment 6 shown in **Figure 3b** was not obtained, indicating that Cas9 cannot simultaneously excise two targeted regions when these are only a few base pairs apart (about 100 bp in our design). This may be due to the Cas9 remaining bound in the cleaved DNA sequence on the 5’-side of the gRNA, allowing preferential ligation of the sequencing adapters to the available end and so providing sequencing directionality (23, 43, 44).

The pool of crRNAs did not cleave the *MYB10*.*3* amplicon tested. Further Sanger sequencing of the uncut amplicon showed that one of the gene copies missed both cleavage target sites. This allele was different from the two *MYB10*.*3* alleles previously identified in ‘Angeleno’, suggesting that the PCR primers had amplified an unknown duplicated *MYB10*.*3* copy. When we later visualized the genome alignments, reads were observed starting from the *MYB10*.*3* annotated position in all the samples, indicating that the Cas9 could indeed cleave this gene with the gRNA pool.

### Sequencing and alignment to reference genomes

The crRNAs were pooled to cleave the DNA of each variety in a single reaction. This strategy made it possible to enclose the ROI and cleave the MYB10 genes along the whole region, regardless of their copy number and organization, which was unknown.

Pooling the barcoded cleaved DNAs lowered the sequencing costs, although it reduced the number of reads obtained per variety, which could have risked the depth of coverage. We obtained a total of 194 Mb of reads, 38.8Mb on average per variety, which represents 288 times the number of bases in the ROI of ‘Sanyueli’, 433 in that of Zhongli-2 and 143 times in Zhongli-1. The depth and coverage obtained after alignment were lower than expected from these figures. The lowest depth values were for the alignments with the ‘Zhongli No. 6’ genome, which was reasonable considering that sequences were aligned to the two assembled MYB10 regions, therefore reducing the depth at each of them. However, the depth in the alignments with ‘Sanyueli’ was only slightly better (from 12.2x to 29.1x, if we dismiss the extremely low values of ‘Golden Japan’). The depth is highly determined by the percentage of coverage and of the on-target reads. We did not observe a correlation between the number of reads obtained and the coverage nor with the on-target ratios. Indeed, the sequencing statistics for ‘Golden Japan’ were similar to those obtained for the other varieties, while all the alignment values were extremely low, indicating a poor enrichment probably caused by an inefficient Cas9 cleavage reaction. The coverage varied per cultivar and reference genome used, although in general it was low (57% on average) due to the low homology between the commercial varieties and *P. salicina* genomes along the ROI, which caused terminal-clipped alignments. The terminal-clipped effect was minor only in the alignment of ‘TC Sun’ - Zhongli-1, which could indicate a shared haplotype between these two cultivars. The ratio of on-target alignments obtained here (8.6%) was higher than reported in other studies (4.61% for multiple cuts in Gilpatrick et al. (23); 2.08 and 3.04% in López-Girona et al. (24). This higher performance could be attributable to the higher number of cleavage sites designed, generating more Cas9-digested DNA fragments with compatible ends for their posterior ligation with the sequencing adapters. This conclusion was also reached by McDonald et al. (38), who obtained a mean on-target value of 44% in the CRISPR Cas9-enrichment of TEs.

Since the complexity of the ROI is out of our control, higher depths would be more easily obtained by increasing the number of cleavage sites. As the crRNAs are pooled in a single cleavage reaction, the cost of the method is only increased by the cost of the crRNAs employed, which is a small proportion of the total cost. In addition, we consider that by reducing the number of pooled varieties, and therefore increasing the number of reads, we would not have obtained much higher coverage or depth. The complexity of the region, its size and the number of cutting sites need to be considered in deciding the number of pooled varieties.

### Variant identification in the alignments to a reference genome

Our results indicate that the strategy described here, applied in a region determining agronomic traits in a panel of selected varieties, can be successfully used to rapidly extract useful information for marker-assisted selection. The generation of long reads offers a higher mapping certainty (45), which is extremely useful for regions with segmental duplications to pinpoint sequence variants with high confidence levels.

We identified SNPs and SVs along the ROI involved in the regulation of fruit skin color. For SNP calling we used Longshot software (46), which not only took into account the error tendency of the ONT long-reads to avoid false positives but has also been reported to outperform the SNP calling method using only short-read sequences in regions with duplications. A total of 129 SNPs were shared by the three cultivars with red-to-purple colored skin and absent in the two with yellow hues; since their phases are known, they could be selected, after validation, as alternatives to the haplotype (PCR) marker provided in Fiol et al. (26).

For SV calling we used Sniffles software (47) complemented with NanoVar (48) to overcome the low depth of ‘Golden Japan’. The accuracy of the pipeline was tested by searching a previously Sanger-validated insertion of 44 bp on the *PsMYB10*.*1a* promoter in the haplotypes associated with red skin color (H1, H3) and in H9. Sniffles could positively detect the polymorphism in homozygosity on ‘Angeleno’ (H1/H3) and in heterozygosity on ‘Black Gold’ (H1/H2), while NanoVar could detect it even in low depth on ‘Golden Japan’ (H4/H9). The insertion was not called in ‘Fortune’ (H3/H6), although some reads were found in low frequency. This indicates that the tools complemented each other well in overcoming their limitations, and that SVs could still be called even in low depth. In addition, we were able to phase the variants, finding the 8 bp insertion in exon 1 of the *PsMYB10*.*1a* gene in the H9 of ‘Golden Japan’ in phase with the 44 bp insertion associated with red skin. This 8 bp insertion produces an early STOP codon, which explains the yellow skin of this cultivar despite it carrying the 44 bp insertion (**Additional File 7**).

### de novo assemblies

We have shown that reads can be aligned to a reference genome or assembled *de novo* in the case of its absence or of bad homology. Here, to recover the positions lost due to the lack of homology with the reference genomes, we used the pipeline for read correction and subsequent *de novo* assembly provided in López-Girona et al. (24). One of the main limitations in our study was that reads produced from two very distant adjacent genes had little chance to overlap. This was the case in the Zhongli-1 region, where two adjacent genes 124 kb apart left 55 kb of the region uncovered. The other limitation for obtaining a whole-region assembly was that reads could not overlap the cleavage positions because these were produced in a single reaction. Therefore, despite reducing the cost, cleaving in a single reaction reduced the accuracy of the assembly, meaning that there should be a compromise between cost and coverage of the assembly when designing the experiment. Due to these limitations, the region was assembled in several contigs, which we put in order using a reference-guided scaffolding method.

The highest sequence conservation and collinearity was between the ‘Angeleno’, ‘Black Gold’ and ‘Fortune’ group, and between the ‘Golden Japan’ and ‘TC Sun’ group comparisons, which was in agreement with the SNP results and the shared haplotypes of the samples. Lower levels of sequence collinearity were observed between samples of the different groups, although generally in several discontinuous blocks of low homology. Given the high complexity of the region and the low conservation levels observed between some haplotypes, we conclude that it was suitable to design crRNAs aimed at the gene sequences, which might be more conserved than the intergenic regions (49, 50).

### Variant identification in the de novo assemblies

The pipeline used to identify variants between *de novo* assemblies succeeded in the identification and assignment of polymorphisms into haplotypes rather than compiling them. The alignments could be separated into haplotypes in three samples and there was enough resolution to correctly identify SNPs and large InDels, improving the results obtained when using a reference genome (the 44 bp insertion was detected in ‘Fortune’). Two samples lacked one of their haplotype sequences, in one case probably due to the high similarity between haplotypes (‘TC Sun’) and in the other due to the low depth of coverage (‘Golden Japan’), therefore, where a reference genome is available, the alignment to it and *de novo* assemblies should be combined to ensure all polymorphisms are identified.

### Strengths, weaknesses, improvements and opportunities

We present here an economic and powerful strategy to extract sequences and polymorphisms from highly complex regions which may contain duplicated and presence/absence variants (PAVs). The main strengths of the strategy are i) that the design of the crRNAs does not necessarily rely on a reference genome, with information of conserved regions or partially cloned alleles being sufficient, ii) that it is economic as the DNA of each variety is cleaved in a single reaction and equimolar amounts of each variety are pooled for a unique ONT run, and iii) that polymorphisms and their phases can be extracted with high efficiency, even in comparison with a reference genome or by analyzing the *de novo* assemblies. The main weaknesses are i) that the search for polymorphisms might require the manual isolation and comparison of contigs (with genes that might have undergone extensive duplication events), ii) that ONT technology is prone to sequence errors (the sequencing errors identified here consist of one and two nucleotide InDels), some of them occurring in short homopolymer regions reported to be an issue in most basecallers (51), and iii) that by using a single Cas9 digestion, overlapping fragments to assemble the region in one contig are not obtained.

This methodology is sufficiently versatile to allow for future improvement. Digesting the DNA region separately with sub-pools of crRNAs would allow for fragment overlapping (this strategy would increase the cost of the experiment, so it is recommended for extremely complicated regions). The scaffolding could be improved by selecting genotypes with the ROI in homozygosis (although this may be difficult to obtain depending on the ROI or the plant materials).

This method offers new opportunities in the identification of genomic variability in agricultural crops, including methylation variants. Changes in methylation patterns may have drastic impacts on the phenotypes. Although we have not explored such application, the methodology used could have allowed it as in Giesselmann et al. (25) and in Gilpatrick et al. (23). In fruit trees, methylation changes in the promoter of *MYB10* genes are associated with changes in fruit color. Also, as in other rosaceous fruit, the fruit color in some Japanese plum varieties is acquired after exposure to cold or light, indicating a possible epigenetic mechanism. Studying methylation changes in the *PsMYB10* region using DNA of the appropriate tissues under certain given conditions may provide important information to understand these regulation mechanisms.

The method that we propose is computationally inexpensive. The most demanding software used in this process is the Flye assembler (52), the cost of which increases with the size of the assembly. In the benchmark tested by the authors, a genome of 4.6 Mb (such as *E. coli)* obtained only with long reads could be assembled using 2 Gb of RAM in about two hours using one CPU. For ROIs less than 1 Mb long it is reasonable to assume that the protocol reported here could be undertaken on most home computers.

## CONCLUSIONS

The CRISPR-Cas9 enrichment strategy enabled long-read sequences from a highly polymorphic and duplicated region to be produced and polymorphisms related to fruit skin color to be deduced from five heterozygous Japanese plum varieties. The methodology was simplified through using a pool of crRNAs in duplicated conserved regions and the sequencing cost was optimized by multiplex sequencing while keeping the protocol computationally inexpensive. The tool could be further improved by taking into account methylation patterns and also adapted for the full assembly of complex regions, being also appropriate for those crops with a lack of a reference genome or with one unrepresentative for the genomic region of interest.

## MATERIALS

### Plant material and nucleic acid isolation

Young leaves from five commercial Japanese plum varieties (**Table 3**) were collected and kept at −80ºC. Prior to DNA extraction, nuclei were separated as described in Naim et al. (53) and detailed online (54), but the vegetal material was homogenized by grinding in liquid nitrogen using a pestle and mortar. DNA was extracted from the isolated nuclei using the Doyle CTAB method (55), introducing an RNAse treatment before the chloroform centrifugation step.

**Table 3.** Commercial varieties selected for CRISPR-Cas9 targeted sequencing of their MYB10 region, their fruit color and the MYB10 haplotype combination described in Fiol et al. (26).

| Variety | Fruit color |  | MYB10 Haplotypes |
| --- | --- | --- | --- |
|  | Skin | Flesh |  |
| ‘Angeleno’ | Black | Yellow | H1/H3 |
| ‘Black Gold’ | Black | Red | H1/H2 |
| ‘Fortune’ | Red | Yellow | H3/H6 |
| ‘Golden Japan’ | Yellow | Yellow | H4/H9 |
| ‘TC Sun’ | Yellow | Yellow | H4/H5 |

The quality of the extracted DNA was evaluated using Nanodrop and low voltage 0.8% agarose TAE gels, and quantified using a Qubit fluorometer.

### Design of guide RNAs targeting conserved sites of *MYB10* genes

The Alt-R CRISPR-Cas9 System (IDT®) was selected to design and assemble the CRISPR RNAs (crRNA) and the trans-activating crRNAs (tracrRNA) into functional guide RNAs (gRNAs).

The crRNA sequences were designed in the desired positions with the IDT custom gRNA design tool. Those with the highest on-target capacity were BLAST to the Peach genome v2.0.a1 (56) and the Sweet cherry genome v1.0 (57) to discard gRNAs with off-target risk regions. SNP variability identified in other genomes was also considered in the crRNA design. Those gRNAs directed at the most highly conserved LG3-MYB10 sequences were finally selected.

### Assembly of the Cas9 RNA-Ribonucleoprotein complex (RNP)

The designed crRNAs and the tracrRNA were resuspended to 100 μM in Nuclease-Free Duplex Buffer (IDT). To form the gRNA duplex, all the crRNAs were pooled in an equimolar mix and combined with the same volume of tracrRNA to a final concentration of 10 μM. The mixture was denaturalized for five minutes at 95ºC and allowed to cool at room temperature. To assemble the Cas9-RNP complex for a single cleavage reaction, 1 μL of the gRNA duplex and 0.08 μL of Alt-R® S.p. HiFi Cas9 Nuclease V3 (IDT) were mixed with 1X of CutSmart Buffer (NEB) and nuclease-free water to a total of 10 μL then incubated for 30 minutes at room temperature and kept on ice for immediate use.

### Cleavage assay of the designed gRNAs with PCR amplicons

The assembled Cas9-RNPs activity was tested using PCR amplicons of *PsMYB10*.*1a, PsMYB10*.*2* and *PsMYB10*.*3* full gene sequences from ‘Angeleno’. The primer sequences and their annealing temperature (Ta) are listed in **Additional File 8**. Each PCR reaction contained 1.5 mM MgCl_2_ and 1X NH_4_ buffers, 1U of BioTaq polymerase (Bioline), 0.2 mM dNTPs, 0.2 μM of each primer, 40 ng of DNA from ‘Angeleno’ and MilliQ water to a total of 25 μL. Thermocycler conditions were 94ºC for 1 minute, 35 cycles of 94ºC for 30 seconds, the Ta described for each primer pair for 20 seconds and 72ºC for 2 minutes, followed by a final extension step of 5 minutes at 72ºC. Reactions were purified using ExoSap-IT (Thermo Fisher) and then 3 μL of each amplicon was diluted with 32 μL of nuclease-free water before adding 10 μL of assembled Cas9-RNP. Cleavage reactions were incubated at 37ºC for 20 minutes and five minutes at 72ºC, then loaded in a 2% agarose TAE gel to visualize the cleaved bands compared to the uncut amplicons.

### Comparison of the MYB10 region in several *Prunus* whole-genome assemblies

A total of fifteen genomes were added to the analysis: two Japanese plums, eight apricots, two sweet cherries, two peaches and a wild peach relative. For all the genomes, the MYB10 region was considered from the BLAST (58) regions of crRNA-f1 and crRNA-f2 sequences. All genomes and the regions considered in this study are listed in **Additional File 2**. The homology between the regions was calculated and visualized pairwise using the DECIPHER v2.0 (59) package for R version 4.0.3 (60). The *MYB10* genes in the two Japanese plum genomes were identified by BLAST using the *PsMYB10* sequences from Fiol et al. (26).

### CRISPR-Cas9 enrichment, library preparation and sequencing

The high-quality nuclei DNA from the five selected commercial plum varieties and the pooled crRNAs (100 μM) were sent to CNAG (Centre Nacional d’Anàlisi Genòmica, Barcelona) for CRISPR-Cas9 enrichment targeted sequencing using Oxford Nanopore Technology (ONT). For each sample, 1000 ng of DNA diluted in 27 μL of nuclease-free water was dephosphorylated by the addition of 3 μL of 10X CutSmart Buffer and 3 μL of Quick Calf Intestinal Phosphatase (NEB) and then incubated at 37ºC for 10 minutes, 80ºC for two minutes and 20ºC for two minutes. To each reaction, 10 μL of previously assembled Cas9-RNP complex was added together with 1 μL of dATP (10 mM) and 1 μL of Taq polymerase (NEB) and incubated at 37ºC for 20 minutes and at 72ºC for five minutes for A-tailing. Samples were tagged using barcodes NB01 to NB05 of the Native Barcoding Expansion 1-12 PCR-free kit (EXP-NBD104, ONT) and purified using Agencourt AMPure XP beads (Beckman Coulter). The barcoded samples were quantified using a Qubit fluorometer and pooled equimolarly to reach 700 ng in 65 μL. The Cas9 Sequencing Kit (SQK-CS9109, ONT) was used to ligate the AMX adapters before purification of the fragments with AMPure XP Beads. The beads were washed twice with Long Fragment Buffer (LFB) and then resuspended in Elution Buffer (EB) for 30 minutes at room temperature. The sequencing library was prepared by mixing 12 μL of the DNA library, 37.5 μL of Sequencing Buffer (SQB) and 25.5 μL of Loading Beads (LB) (SQK-CS9109, ONT). The sample of pooled fragments was sequenced in a MinION (v9.4.1) flowcell on a GridION mK1 device operated by MinKNOW version 3.6.5 software.

### Basecalling, quality filtering, adapter removal, alignment to reference and variant calling

Nanopore read sequences were basecalled and demultiplexed using Guppy (v3.2.10+aabd4ec) high accuracy model (ONT). MinIONQC v1.4.2 (61) was used to evaluate the read quality scores before and after trimming barcodes and adapters with Porechop v0.2.4 (https://github.com/rrwick/Porechop). Reads were aligned to *Prunus salicina* ‘Sanyueli’ v1.0 (27) and *P. salicina* ‘Zhongli No. 6’ v1.0 (28) whole genome assemblies by using Minimap2 (62), and were visualized in IGV (63) and represented in bwtool (64). Alignment information was obtained with Samtools version 1.11 (65). Variant calling was done with software specifically scripted for third-generation sequencing reads: Longshot v0.4.3 (46) was run for SNP calling, while both Sniffles v1.0.12 (47) and NanoVar v1.3.9 (48) were run, and their output files merged with VCFtools v0.1.16 (66), to obtain the list of SV calls. VFCtools was further used to compare the lists of SNP calls between the five ‘Sanyueli’ genome alignments, and the results represented in a Venn diagram using pyvenn (https://github.com/tctianchi/pyvenn).

### Reads correction, Flye assembly and reference-guided scaffolding

Canu v 2.1.1 (67) was used to correct the trimmed reads and perform an initial *de novo* assembly. Nanopolish v0.11.1 (68) was used to improve the assembly results. For each sequenced cultivar, their Canu corrected reads and polished contigs were used as input for Flye assembler v2.8.3 (52) enabling the option to keep haplotypes. The resulting contigs were genome-guided scaffolded with the RagTag software (69), using separately the three MYB10 regions from the two *P. salicina* available genomes as template reference sequences. DECIPHER v2.0 (59) was used to display the homology between the references and reference-guided sequences in a pairwise fashion and obtain their similarity values.

### Identification of SNPs and InDels in the *PsMYB10*.*1a* promoter

A region of 250 bp in the *PsMYB10*.*1a* promoter containing several SNPs and large InDels was used for comparison. Sanger sequences of this region obtained for different haplotypes

(26) were used in a BLAST analysis to recover the haplotypes in the *de novo* (Flye) contigs. In the case of the H4 sequence, which could not be recovered, reads were isolated from their alignment to the ‘Zhongli No. 6’ genome *PsMYB10*.*1a* promoter region and assembled manually. No Sanger sequences were available for H5 and H6, and their contigs on ‘TC Sun’ and ‘Fortune’ were isolated as alternative to H4 and H3 sequences. The selected contigs from each cultivar were aligned and compared using Sequencher 5.0 (Gene Codes Corporation, Ann Arbor, MI, USA).

## Availability of data and materials

The raw sequencing data presented in this study has been deposited at the EBI European Nucleotide Archive repository under the accession number PRJEB48338.

## Authors’ contributions

The experiments were conceived and designed by AF and MJA. ELG provided support in the design of the experiment. Experiments were conducted by AF. Bioinformatics analysis were performed by AF and FJR. The paper was written by AF and MJA. All authors have critically revised the manuscript and approved the final document.

## Acknowledgements

We thank Joan París, Ferran Contreras (ADV de Fruita del Baix Llobregat) and Jorge Naranjo (Tany Nature) for facilitating the plant material used in the experiment. We would like to thank David Chagne for thoughtful comments and helpful advice while reviewing our manuscript.

## Additional files

**Additional File 1**. Dot plot comparing, pair-wise, the MYB10 regions identified in 15 *Prunus* genomes, represented as in Figure 1a. The colored squares border the *Prunus* sections considered: purple for Japanese plums, orange for apricots, red for sweet cherries, and green for peaches and its wild relative.

**Additional File 2**. Details of the *MYB10* region isolated from each of the fifteen *Prunus* genomes considered for pair-wise comparison. The results of the percentage of sequence hits and the number of shared blocks for every comparison is included.

**Additional File 3**. Details of the crRNAs designed for the Japanese plum LG3-MYB10 region enrichment. The SNPs identified are underlined and crRNAs were designed including each variant.

**Additional File 4**. Visualization of the depth of the sequences aligned to the ‘Sanyueli’, Zhongli-1 and Zhongli-2 regions. The pink bars highlight the coordinates with previously identified *MYB10* gene sequences, where the Cas9 enzyme has two cutting points enabling sequencing in both directions.

**Additional File 5**. The total number of SNPs called from each sequenced sample and the count of the shared SNP positions between them.

**Additional File 6**. Pair-wise visualization of the homologous hits (above diagonal) and homologous blocks (below) between each reference region and the *de novo* contigs scaffolded using the ‘Sanyueli’, Zhongli-1 or Zhongli-2 regions, represented in the same colors as in Figure 1a.

**Additional File 7**. Visualization of two phased variants 1 kb apart on H9. The 44 bp insertion is present in H1, H3 and H9 and was associated to the red skin color. The polymorphism is phased with an 8 bp insertion at the start of exon 1 of the *PsMYB10*.*1* gene, which explains its lack of function on H9. The reads from H4 do not show either of the two polymorphisms.

**Additional File 8**. Primer sequences and their annealing temperatures (Ta) used to PCR amplify the *MYB10* gene sequences.

## References

1. Taylor JS, Raes J. Duplication and divergence: the evolution of new genes and old ideas. Annu Rev Genet. 2004;38:615–43.

2. Van de Peer Y, Maere S, Meyer A. The evolutionary significance of ancient genome duplications. Nat Rev Genet. 2009;10(10):725–32.

3. Gabur I, Chawla HS, Snowdon RJ, Parkin IA. Connecting genome structural variation with complex traits in crop plants. Theor Appl Genet. 2019;132(3):733–50.

4. Soltis DE, Visger CJ, Soltis PS. The polyploidy revolution then…and now: Stebbins revisited. Am J Bot. 2014;101(7):1057–78.

5. Giannuzzi G, D’Addabbo P, Gasparro M, Martinelli M, Carelli FN, Antonacci D, et al. Analysis of high-identity segmental duplications in the grapevine genome. BMC Genomics. 2011;12(1):1–14.

6. Fares MA, Keane OM, Toft C, Carretero-Paulet L, Jones GW. The roles of whole-genome and small-scale duplications in the functional specialization of Saccharomyces cerevisiae genes. Plos Genet. 2013;9(1):e1003176.

7. Zmienko A, Marszalek-Zenczak M, Wojciechowski P, Samelak-Czajka A, Luczak M, Kozlowski P, et al. AthCNV: A map of DNA copy number variations in the Arabidopsis genome. The Plant Cell. 2020;32(6):1797–819.

8. Vandepoele K, Simillion C, Van de Peer Y. Evidence that rice and other cereals are ancient aneuploids. The Plant Cell. 2003;15(9):2192–202.

9. Lye ZN, Purugganan MD. Copy number variation in domestication. Trends Plant Sci. 2019;24(4):352–65.

10. Xu C, Nadon BD, Kim KD, Jackson SA. Genetic and epigenetic divergence of duplicate genes in two legume species. Plant, cell & environment. 2018;41(9):2033–44.

11. Gu YQ, Crossman C, Kong X, Luo M, You FM, Coleman-Derr D, et al. Genomic organization of the complex α-gliadin gene loci in wheat. Theor Appl Genet. 2004;109(3):648–57.

12. Camerlengo F, Sestili F, Silvestri M, Colaprico G, Margiotta B, Ruggeri R, et al. Production and molecular characterization of bread wheat lines with reduced amount of α-type gliadins. BMC Plant Biol. 2017;17(1):1–11.

13. Ruiz M, Giraldo P, Royo C, Villegas D, Aranzana MJ, Carrillo JM. Diversity and Genetic Structure of a Collection of Spanish Durum Wheat Landraces. Crop Sci. 2012;52(5):2262–75.

14. Van Herpen TW, Goryunova SV, Van Der Schoot J, Mitreva M, Salentijn E, Vorst O, et al. Alpha-gliadin genes from the A, B, and D genomes of wheat contain different sets of celiac disease epitopes. BMC Genomics. 2006;7(1):1–13.

15. Chawla HS, Lee H, Gabur I, Vollrath P, Tamilselvan-Nattar-Amutha S, Obermeier C, et al. Long-read sequencing reveals widespread intragenic structural variants in a recent allopolyploid crop plant. Plant Biotechnol J. 2021;19(2):240–50.

16. Alonge M, Wang X, Benoit M, Soyk S, Pereira L, Zhang L, et al. Major impacts of widespread structural variation on gene expression and crop improvement in tomato. Cell. 2020;182(1):145-61. e23.

17. Della Coletta R, Qiu Y, Ou S, Hufford MB, Hirsch CN. How the pan-genome is changing crop genomics and improvement. Genome Biol. 2021;22(1):1–19.

18. Tao Y, Zhao X, Mace E, Henry R, Jordan D. Exploring and exploiting pan-genomics for crop improvement. Mol Plant. 2019;12(2):156–69.

19. She X, Jiang Z, Clark RA, Liu G, Cheng Z, Tuzun E, et al. Shotgun sequence assembly and recent segmental duplications within the human genome. Cah Rev The. 2004;431(7011):927–30.

20. Amarasinghe SL, Su S, Dong X, Zappia L, Ritchie ME, Gouil Q. Opportunities and challenges in long-read sequencing data analysis. Genome Biol. 2020;21(1):1–16.

21. Liu Y, Du H, Li P, Shen Y, Peng H, Liu S, et al. Pan-genome of wild and cultivated soybeans. Cell. 2020;182(1):162-76.e13.

22. Song J-M, Guan Z, Hu J, Guo C, Yang Z, Wang S, et al. Eight high-quality genomes reveal pan-genome architecture and ecotype differentiation of Brassica napus. Nature Plants. 2020;6(1):34–45.

23. Gilpatrick T, Lee I, Graham JE, Raimondeau E, Bowen R, Heron A, et al. Targeted nanopore sequencing with Cas9-guided adapter ligation. Nat Biotechnol. 2020;38(4):433–8.

24. López-Girona E, Davy MW, Albert NW, Hilario E, Smart ME, Kirk C, et al. CRISPR-Cas9 enrichment and long read sequencing for fine mapping in plants. Plant Methods. 2020;16(1):1–13.

25. Giesselmann P, Brändl B, Raimondeau E, Bowen R, Rohrandt C, Tandon R, et al. Analysis of short tandem repeat expansions and their methylation state with nanopore sequencing. Nat Biotechnol. 2019;37(12):1478–81.

26. Fiol A, García-Gómez BE, Jurado-Ruiz F, Alexiou K, Howad W, Aranzana MJ. Characterization of Japanese Plum (Prunus salicina) PsMYB10 Alleles Reveals Structural Variation and Polymorphisms Correlating With Fruit Skin Color. Front Plant Sci. 2021;12:1057.

27. Liu C, Feng C, Peng W, Hao J, Wang J, Pan J, et al. Chromosome-level draft genome of a diploid plum (Prunus salicina). GigaScience. 2020;9(12):giaa130.

28. Huang Z, Shen F, Chen Y, Cao K, Wang L. Chromosome-scale genome assembly and population genomics provide insights into the adaptation, domestication, and flavonoid metabolism of Chinese plum. Plant J. 2021.

29. Panchy N, Lehti-Shiu M, Shiu S-H. Evolution of gene duplication in plants. Plant Physiol. 2016;171(4):2294–316.

30. González VM, Aventín N, Centeno E, Puigdomènech P. High presence/absence gene variability in defense-related gene clusters of Cucumis melo. BMC Genomics. 2013;14(1):1–13.

31. Qiao X, Yin H, Li L, Wang R, Wu J, Wu J, et al. Different modes of gene duplication show divergent evolutionary patterns and contribute differently to the expansion of gene families involved in important fruit traits in pear (Pyrus bretschneideri). Front Plant Sci. 2018;9:161.

32. Gu C, Wang L, Wang W, Zhou H, Ma B, Zheng H, et al. Copy number variation of a gene cluster encoding endopolygalacturonase mediates flesh texture and stone adhesion in peach. J Exp Bot. 2016;67(6):1993–2005.

33. Barragan AC, Weigel D. Plant NLR diversity: the known unknowns of Pan-NLRomes. The Plant Cell. 2021;33(4):814–31.

34. Badet T, Croll D. The rise and fall of genes: origins and functions of plant pathogen pangenomes. Curr Opin Plant Biol. 2020;56:65–73.

35. Lallemand T, Leduc M, Landès C, Rizzon C, Lerat E. An overview of duplicated gene detection methods: Why the duplication mechanism has to be accounted for in their choice. Genes. 2020;11(9):1046.

36. Gabrieli T, Sharim H, Fridman D, Arbib N, Michaeli Y, Ebenstein Y. Selective nanopore sequencing of human BRCA1 by Cas9-assisted targeting of chromosome segments (CATCH). Nucleic Acids Res. 2018;46(14):e87–e.

37. Bruijnesteijn J, van der Wiel M, de Groot NG, Bontrop RE. Rapid characterization of complex genomic regions using Cas9 enrichment and Nanopore sequencing. bioRxiv. 2021.

38. McDonald TL, Zhou W, Castro CP, Mumm C, Switzenberg JA, Mills RE, et al. Cas9 targeted enrichment of mobile elements using nanopore sequencing. Nat Commun. 2021;12(1):1–13.

39. Kirov I, Merkulov P, Gvaramiya S, Komakhin R, Omarov M, Dudnikov M, et al. Illuminating the transposon insertion landscape in plants using Cas9-targeted Nanopore sequencing and a novel pipeline. bioRxiv. 2021.

40. Okie W, Ramming D. Plum breeding worldwide. Horttechnology. 1999;9(2):162–76.

41. Jung S, Jiwan D, Cho I, Lee T, Abbot AG, Sosinski B, et al. Synteny of Prunus and other model plant species. BMC Genomics. 2009;10:76.

42. Stracke R, Werber M, Weisshaar B. The R2R3-MYB gene family in Arabidopsis thaliana. Curr Opin Plant Biol. 2001;4(5):447–56.

43. Sternberg SH, Redding S, Jinek M, Greene EC, Doudna JA. DNA interrogation by the CRISPR RNA-guided endonuclease Cas9. Cah Rev The. 2014;507(7490):62–7.

44. Brinkman EK, Chen T, de Haas M, Holland HA, Akhtar W, van Steensel B. Kinetics and fidelity of the repair of Cas9-induced double-strand DNA breaks. Mol Cell. 2018;70(5):801-13.e6.

45. Li W, Freudenberg J. Mappability and read length. Frontiers in genetics. 2014;5:381.

46. Edge P, Bansal V. Longshot enables accurate variant calling in diploid genomes from single-molecule long read sequencing. Nat Commun. 2019;10(1):1–10.

47. Sedlazeck FJ, Rescheneder P, Smolka M, Fang H, Nattestad M, Von Haeseler A, et al. Accurate detection of complex structural variations using single-molecule sequencing. Nat Methods. 2018;15(6):461–8.

48. Tham CY, Tirado-Magallanes R, Goh Y, Fullwood MJ, Koh BT, Wang W, et al. NanoVar: accurate characterization of patients’ genomic structural variants using low-depth nanopore sequencing. Genome Biol. 2020;21(1):1–15.

49. Lynch M, Force A. The probability of duplicate gene preservation by subfunctionalization. Genetics. 2000;154(1):459–73.

50. Kondrashov FA, Rogozin IB, Wolf YI, Koonin EV. Selection in the evolution of gene duplications. Genome Biol. 2002;3(2):1–9.

51. Wick RR, Judd LM, Holt KE. Performance of neural network basecalling tools for Oxford Nanopore sequencing. Genome Biol. 2019;20(1):1–10.

52. Kolmogorov M, Yuan J, Lin Y, Pevzner PA. Assembly of long, error-prone reads using repeat graphs. Nat Biotechnol. 2019;37(5):540–6.

53. Naim F, Nakasugi K, Crowhurst RN, Hilario E, Zwart AB, Hellens RP, et al. Advanced engineering of lipid metabolism in Nicotiana benthamiana using a draft genome and the V2 viral silencing-suppressor protein. Plos One. 2012;7(12):e52717.

54. Hilario E. Plant nuclear genomic DNA preps 2018 [Available from: https://www.protocols.io/view/plant-nuclear-genomic-dna-preps-rncd5aw.

55. Doyle JJ, Doyle JL. A rapid DNA isolation procedure for small quantities of fresh leaf tissue. 1987.

56. Verde I, Jenkins J, Dondini L, Micali S, Pagliarani G, Vendramin E, et al. The Peach v2.0 release: high-resolution linkage mapping and deep resequencing improve chromosome-scale assembly and contiguity. BMC Genomics. 2017;18(1):225.

57. Shirasawa K, Isuzugawa K, Ikenaga M, Saito Y, Yamamoto T, Hirakawa H, et al. The genome sequence of sweet cherry (Prunus avium) for use in genomics-assisted breeding. DNA Res. 2017;24(5):499–508.

58. Altschul SF, Gish W, Miller W, Myers EW, Lipman DJ. Basic local alignment search tool. J Mol Biol. 1990;215(3):403–10.

59. Wright ES. Using DECIPHER v2. 0 to analyze big biological sequence data in R. R Journal. 2016;8(1).

60. R Core Team. R: A language and environment for statistical computing. R Foundation for Statistical Computing. Vienna, Austria.: URL http://www.R-project.org/. 2020.

61. Lanfear R, Schalamun M, Kainer D, Wang W, Schwessinger B. MinIONQC: fast and simple quality control for MinION sequencing data. Bioinformatics. 2019;35(3):523–5.

62. Li H. Minimap2: pairwise alignment for nucleotide sequences. Bioinformatics. 2018;34(18):3094–100.

63. Robinson JT, Thorvaldsdóttir H, Winckler W, Guttman M, Lander ES, Getz G, et al. Integrative genomics viewer. Nat Biotechnol. 2011;29(1):24–6.

64. Pohl A, Beato M. bwtool: a tool for bigWig files. Bioinformatics. 2014;30(11):1618–9.

65. Li H, Handsaker B, Wysoker A, Fennell T, Ruan J, Homer N, et al. The sequence alignment/map format and SAMtools. Bioinformatics. 2009;25(16):2078–9.

66. Danecek P, Auton A, Abecasis G, Albers CA, Banks E, DePristo MA, et al. The variant call format and VCFtools. Bioinformatics. 2011;27(15):2156–8.

67. Koren S, Walenz BP, Berlin K, Miller JR, Bergman NH, Phillippy AM. Canu: scalable and accurate long-read assembly via adaptive k-mer weighting and repeat separation. Biotechfor. 2017;27(5):722–36.

68. Loman NJ, Quick J, Simpson JT. A complete bacterial genome assembled de novo using only nanopore sequencing data. Nat Methods. 2015;12(8):733–5.

69. Alonge M, Soyk S, Ramakrishnan S, Wang X, Goodwin S, Sedlazeck FJ, et al. RaGOO: fast and accurate reference-guided scaffolding of draft genomes. Genome Biol. 2019;20(1):1–17.

