## Supplementary figures and images for "An efficient CRISPR-Cas9 enrichment sequencing strategy for characterizing complex and highly duplicated genomic regions. A case study in the *Prunus salicina* LG3-MYB10 genes cluster"

### Additional File 1

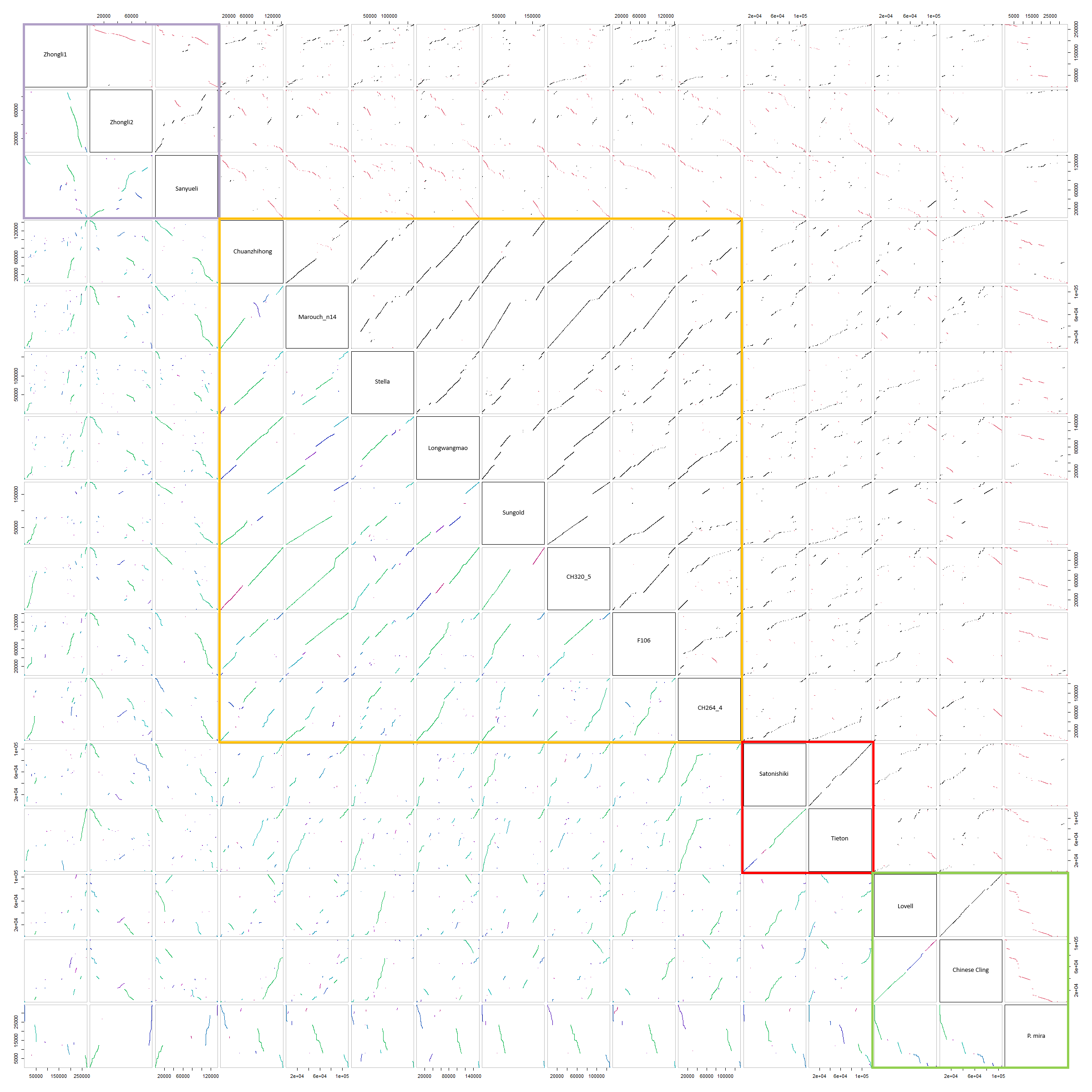

### Additional File 7

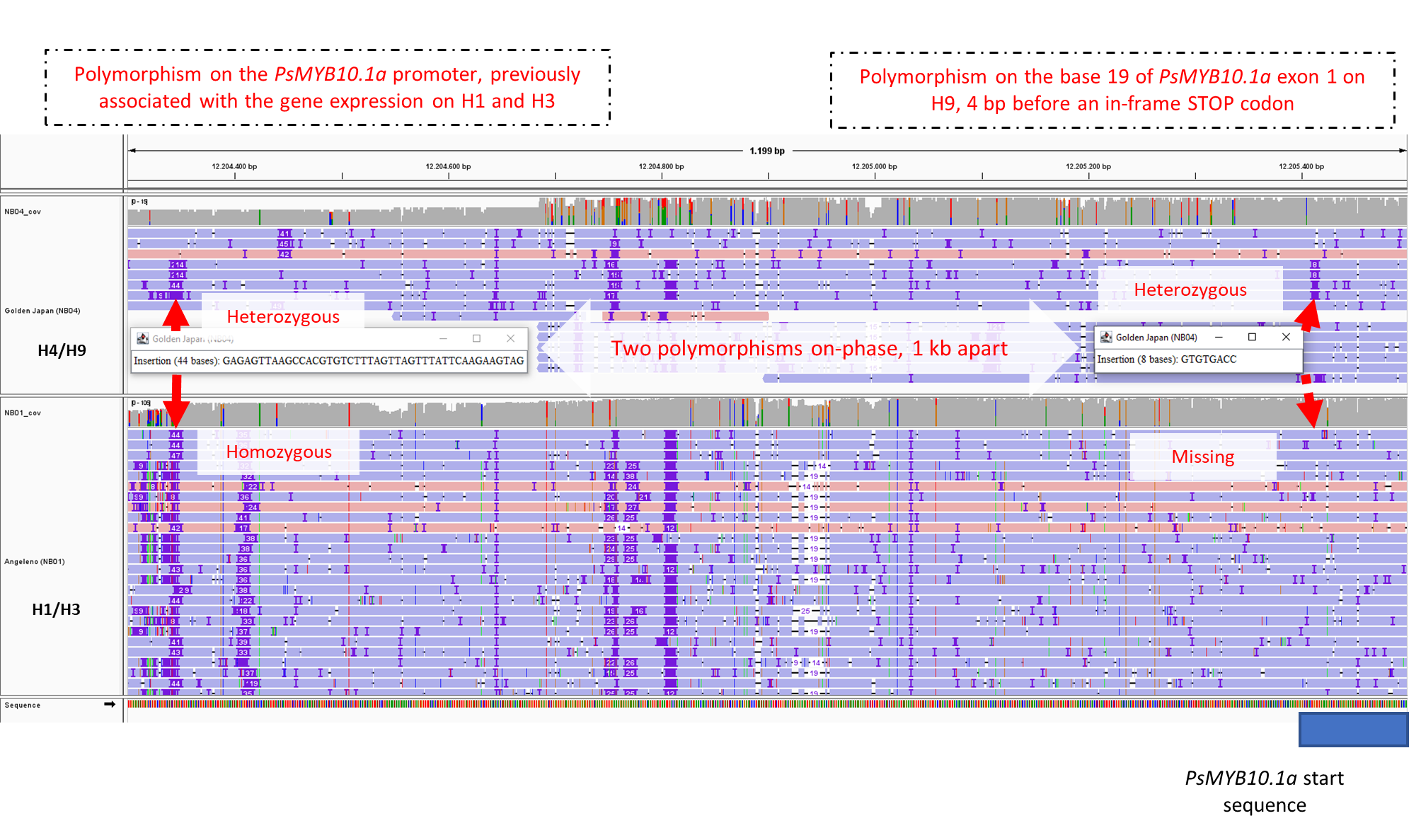
