## Additional File 3 for "An efficient CRISPR-Cas9 enrichment sequencing strategy for characterizing complex and highly duplicated genomic regions. A case study in the *Prunus salicina* LG3-MYB10 genes cluster"

**Additional File 3.** Details of the crRNAs designed for Japanese plum LG3-MYB10 region enrichment. The SNPs identified are shown underlined and crRNAs were designed including each variant.

| **Name** | **Sequence 5’->3’** | **PAM** | **Target** | **Strand** | **On-target activity score** |
| --- | --- | --- | --- | --- | --- |
| **crRNA-s1** | GGAAGAGCTGTAGACTAAGG | TGG | MYB10.1 and MYB10.2 (exon 2) | (+) | 58 |
| **crRNA-s2** | GGAGGAGCTGTAGACTAAGG | TGG | MYB10.2 (exon 2) | (+) | 50 |
| **crRNA-s3** | GGAAGAGCTGCAGACTACGG | TGG | MYB10.3 (exon 2) | (+) | 57 |
| **crRNA-a1** | ATAAGTCTCTTAGCACCCCT | CGG | MYB10.1 (intron 1) | (-) | 64 |
| **crRNA-a2** | ATAAGTCTCTTAGTACCCCT | CGG | MYB10.1 (intron 1) | (-) | 67 |
| **crRNA-a3** | AAACTTGTCATGAAATTATC | AGG | MYB10.2 (intron 1) | (-) | 51 |
| **crRNA-a4** | ATTGTATAACATCTTTCTCG | AGG | MYB10.3 (intron 1) | (-) | 68 |
| **crRNA-f1** | CGGGTGCAAGGCGTACCAAG | CGG | *Prupe.3G162900* | (-) | 70 |
| **crRNA-f2** | GACTACCCCTGCGAGTCCAG | AGG | *Prupe.3G163400* | (+) | 61 |
