## Additional File 4 for "An efficient CRISPR-Cas9 enrichment sequencing strategy for characterizing complex and highly duplicated genomic regions. A case study in the *Prunus salicina* LG3-MYB10 genes cluster"

### Slide 1
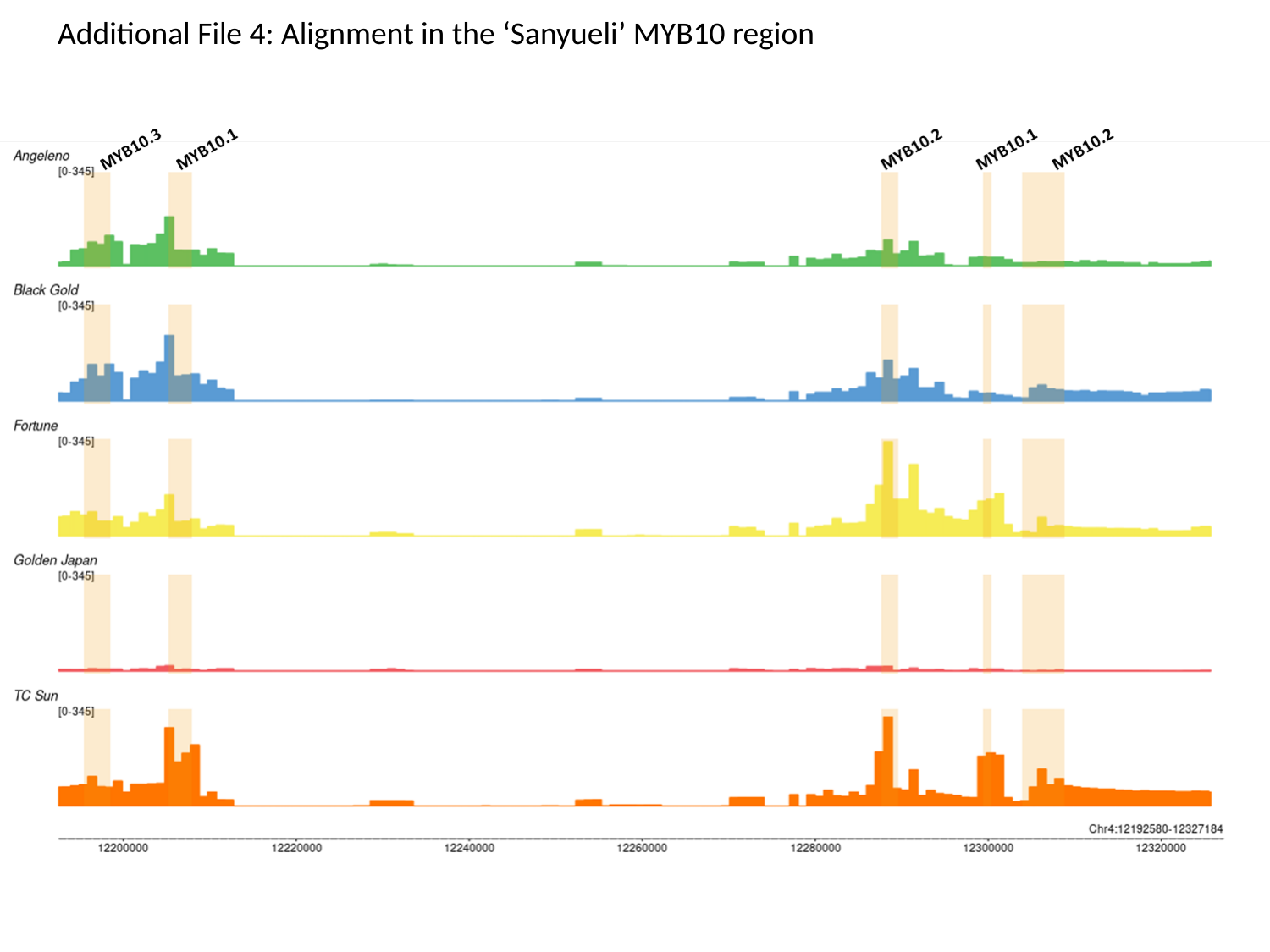

Additional File 4: Alignment in the ‘Sanyueli’ MYB10 region

### Slide 2
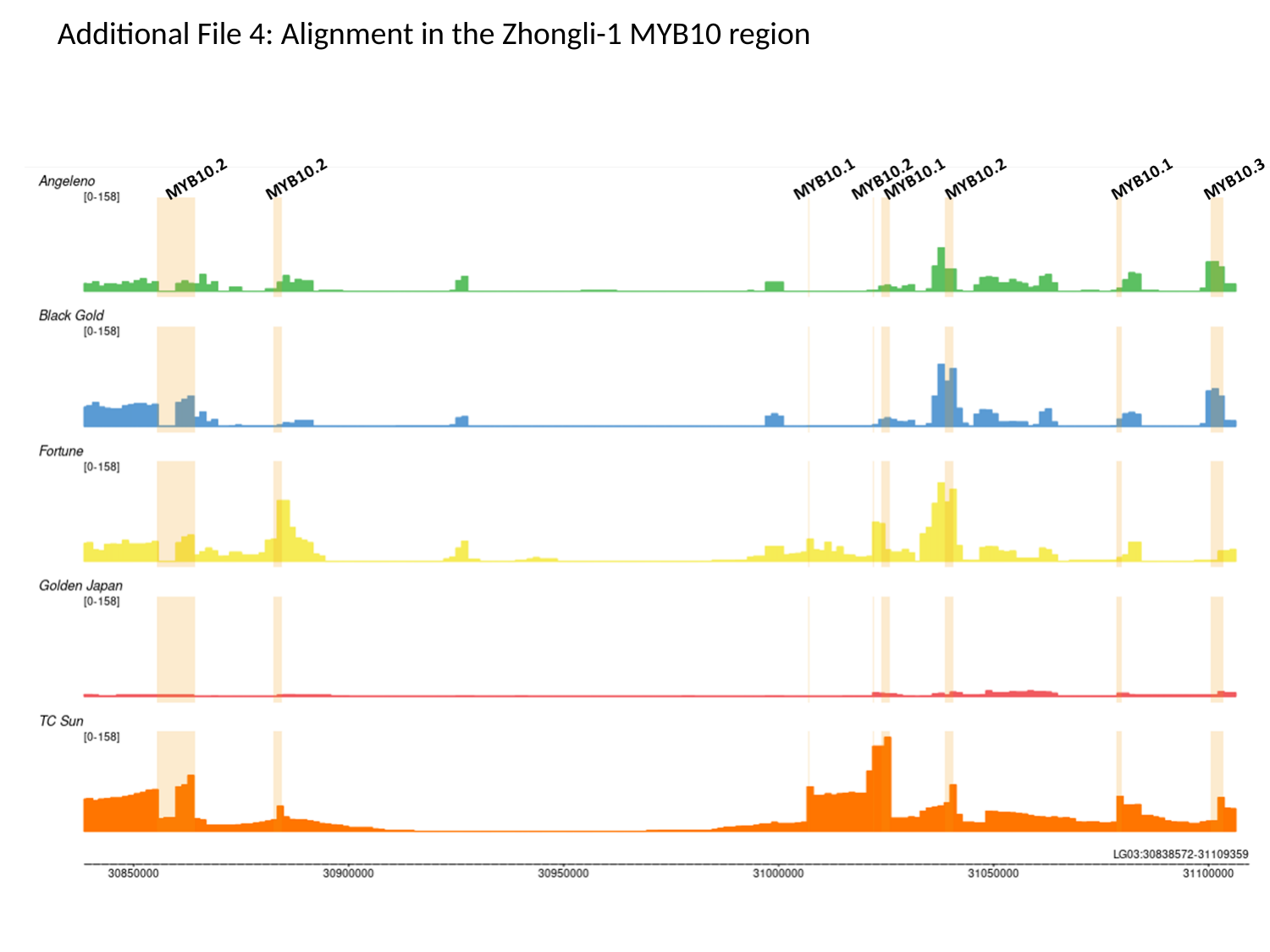

Additional File 4: Alignment in the Zhongli-1 MYB10 region

### Slide 3
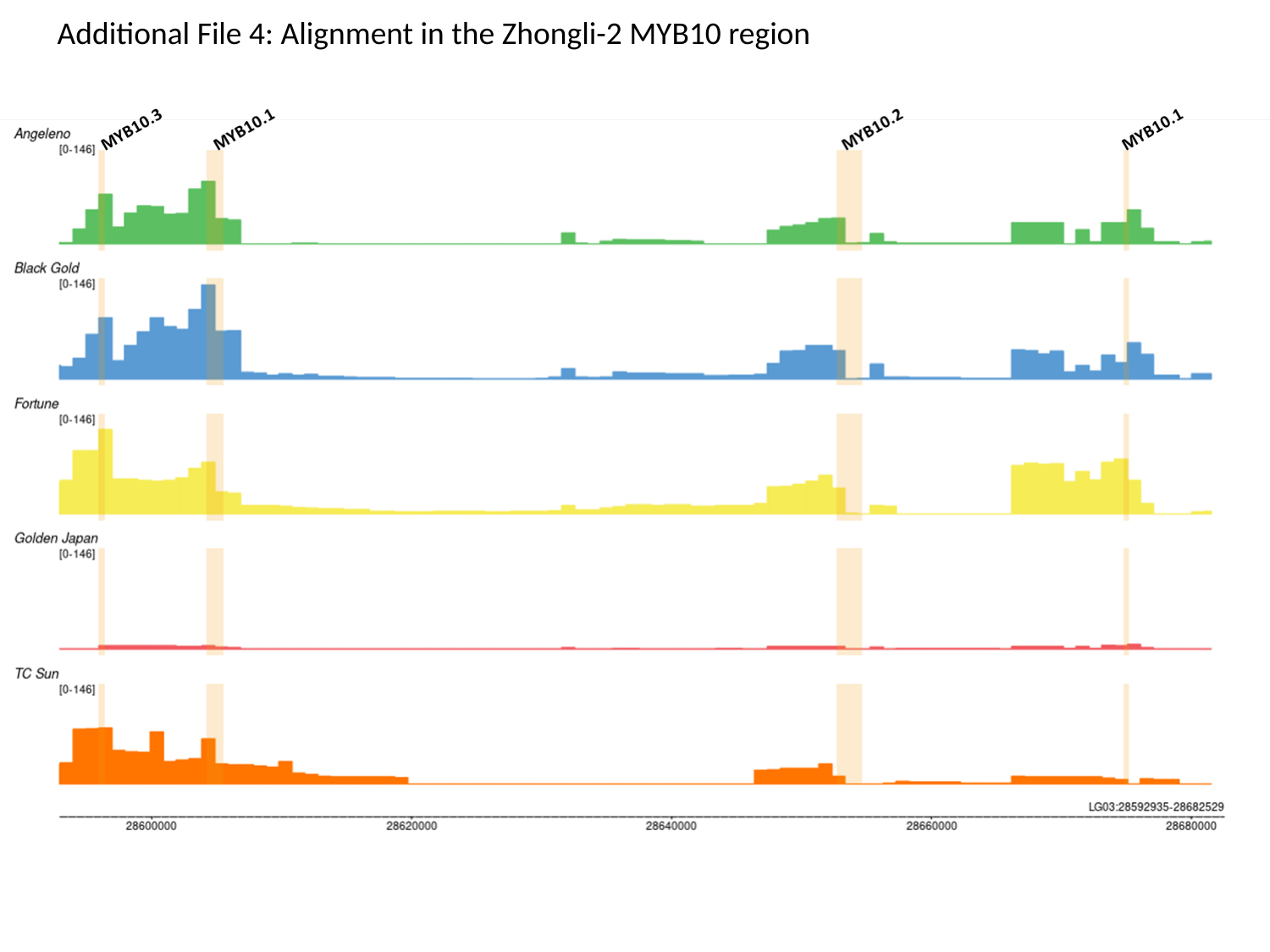

Additional File 4: Alignment in the Zhongli-2 MYB10 region
