## Additional File 6 for "An efficient CRISPR-Cas9 enrichment sequencing strategy for characterizing complex and highly duplicated genomic regions. A case study in the *Prunus salicina* LG3-MYB10 genes cluster"

### Slide 1
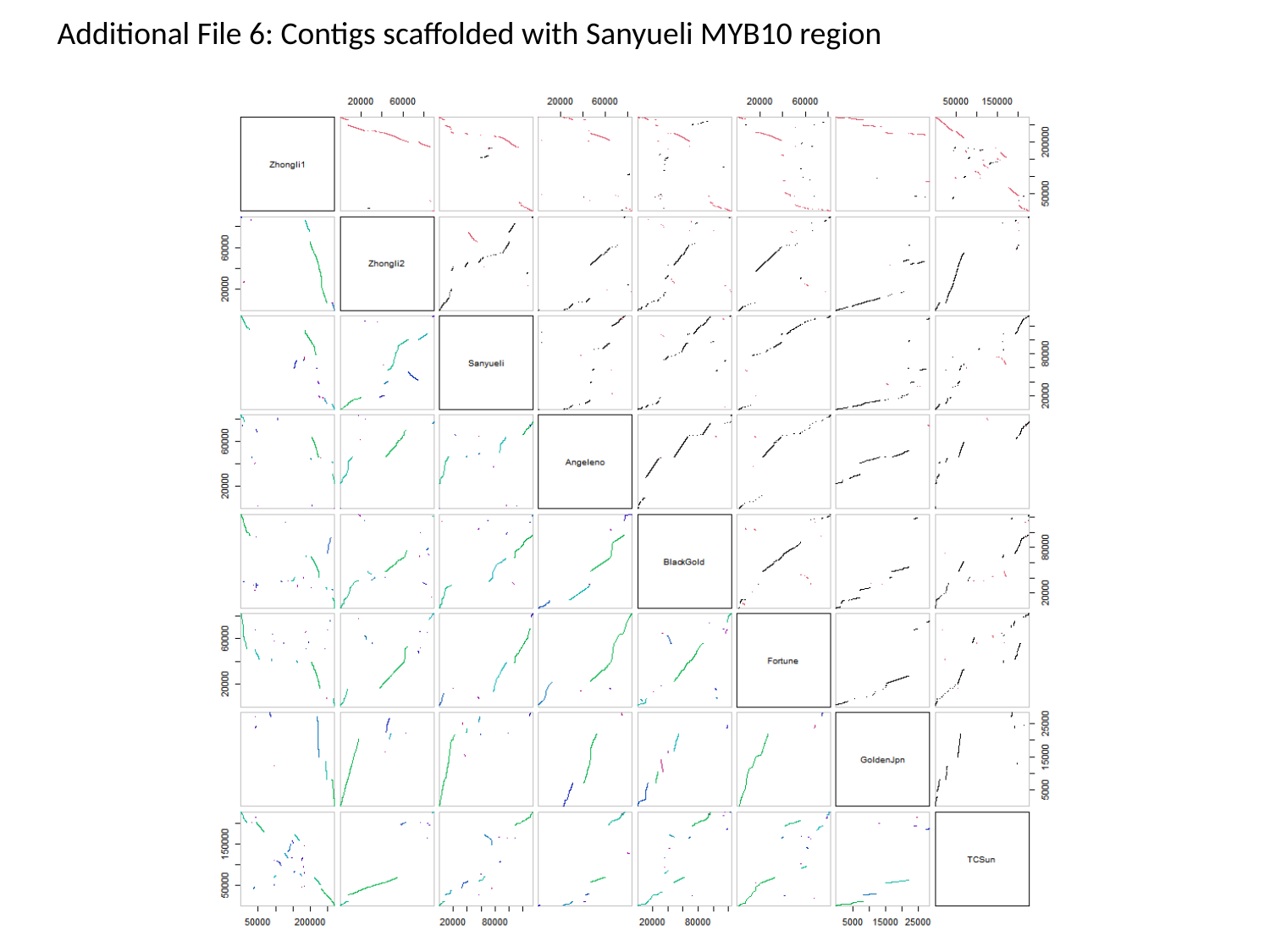

Additional File 6: Contigs scaffolded with Sanyueli MYB10 region

### Slide 2
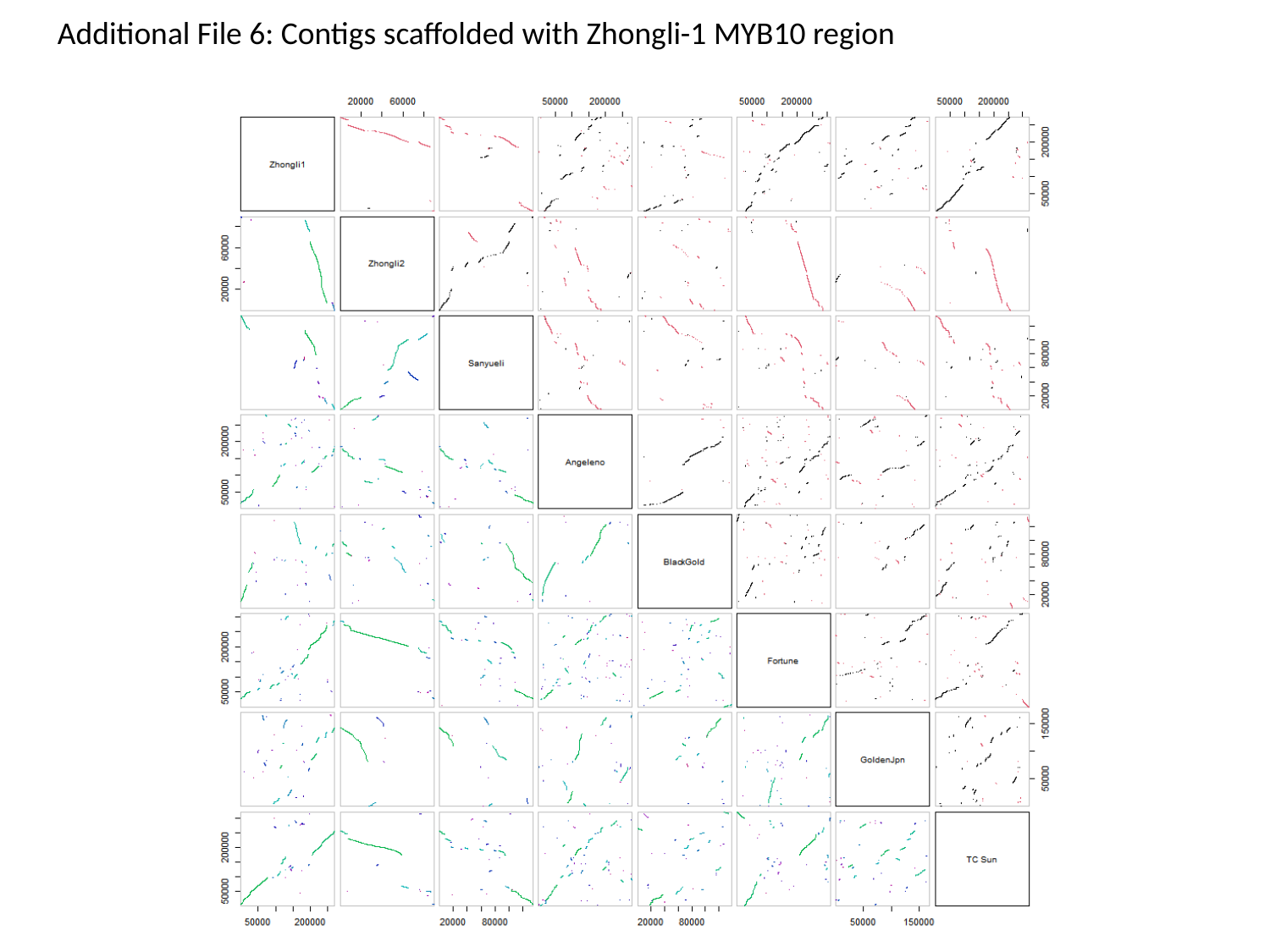

Additional File 6: Contigs scaffolded with Zhongli-1 MYB10 region

### Slide 3
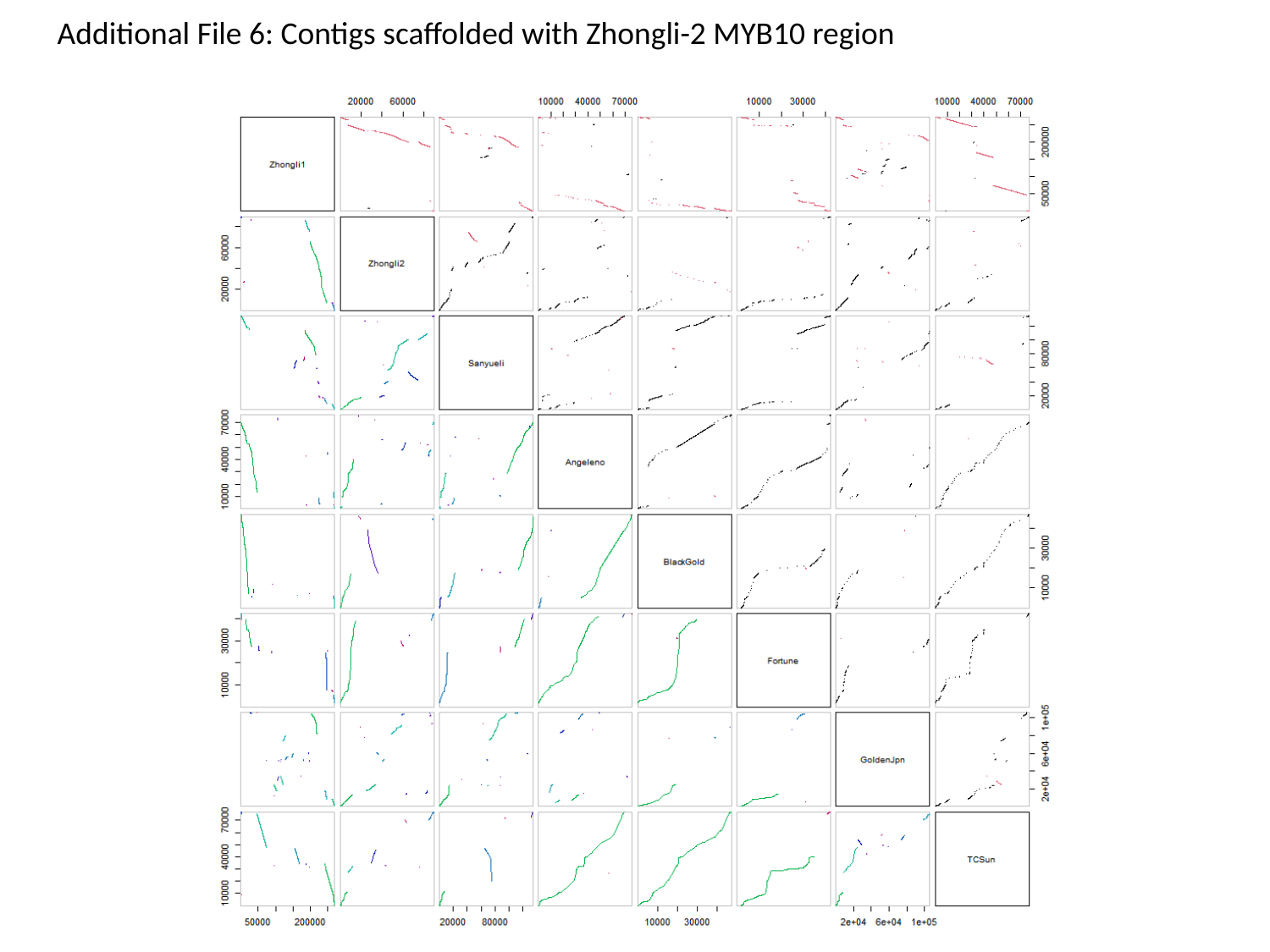

Additional File 6: Contigs scaffolded with Zhongli-2 MYB10 region
