## Additional File 8 for "An efficient CRISPR-Cas9 enrichment sequencing strategy for characterizing complex and highly duplicated genomic regions. A case study in the *Prunus salicina* LG3-MYB10 genes cluster"

| **Primer name** | **Sequence** | **Ta (ºC)** | **Description** |
| --- | --- | --- | --- |
| **M101_RT_F** | TGGACACGAGACATTGCACG | 57 | *PsMYB10.1* amplification (Fiol et al., 2021) |
| **M101_RT_R** | CAATGGTCTTTTGACAGCCGC |  |  |
| **M102_f** | CTGGCTGCAAGCATAC | 57 | *PsMYB10.2* amplification (Fiol et al., 2021) |
| **M102_r** | GTGGGACAAACACTCTC |  |  |
| **M103_f** | ATAGGAACTAGCAGGCAC | 57 | *PsMYB10.3* amplification (Fiol et al., 2021) |
| **M103_r** | AGTTGCTAATAATTGCTACTAGG |  |  |

**Additional File 8.** Primer sequences and their temperatures of annealing (Ta) used to PCR amplify the *MYB10* gene sequences.
